# Dynamic Meta-Networking Identifies Distinct Network Correlates of Positive and Negative Formal Thought Disorder in Schizophrenia

**DOI:** 10.1101/2025.09.29.679140

**Authors:** Zhe Hu, Jianshan Chen, Junjie Yang, Hui Xie, Jiaxuan Liu, Peifen Zhang, Qiuxia Wu, Mohammad-Hossein Doosti, Cathy Rong, Jijun Wang, Xueqin Song, Xiaochen Tang, Binke Yuan, Liping Cao

## Abstract

Formal thought disorder (FTD) is a core symptom of schizophrenia, yet the neural network mechanisms underlying this phenotype remain poorly understood. In this study, we applied a dynamic meta-networking framework, which captures temporally recurring functional network states, to investigate alterations in the language and executive control networks and their associations with positive and negative FTD. Resting-state fMRI data were collected from three independent cohorts: a discovery cohort comprising 150 first-episode, drug-naïve patients with schizophrenia and 175 healthy controls (HCs); a replication cohort including 183 first-episode, drug-naïve patients and 109 HCs; and a third cohort consisting of 71 patients who had received two weeks of antipsychotic treatment and 71 HCs. Meta-networking analysis identified four distinct resting-state meta-states within both the language and executive control networks. Connectivity-behavior correlation analyses and machine-learning regression models revealed that positive FTD was associated with aberrant connectivity across specific meta-states in both networks. In contrast, negative FTD was linked exclusively to dysfunction within two meta-states of the executive control network. Notably, these polarity-specific, multi-state connectivity disruptions normalized following short-term antipsychotic treatment, highlighting their potential as clinically relevant neuroimaging biomarkers.

## Introduction

Formal thought disorder (FTD) is clinically recognized as a disruption of language, thinking, and communication processes^1,2^. FTD is a core feature of schizophrenia (SZ), often emerging during the prodromal phase and persisting throughout the disease trajectory^3^, significantly impairing communication and functional outcomes. Understanding the neural mechanisms underlying FTD is critical for elucidating schizophrenia pathophysiology and for identifying biomarkers that could enable early detection and disease monitoring^4^. Standardized clinical scales, including the Scale for the Assessment of Positive Symptoms (SAPS), the Scale for the Assessment of Negative Symptoms (SANS), and the Positive and Negative Syndrome Scale (PANSS), classify FTD into positive and negative dimensions. Positive FTD is characterized by disorganized speech and thought, manifesting as derailment, tangentiality, illogicality, and peculiar speech patterns. In contrast, negative FTD is characterized by diminished thought and speech activity, often resulting in impoverished, content-deficient communication^2,5^. The distinct symptomatology suggests the involvement of potentially divergent underlying pathophysiological mechanisms^6^.

Two prominent frameworks have emerged to explain the neurocognitive basis of FTD: the dyssemantic and dysexecutive hypotheses^6–8^. The dyssemantic hypothesis proposes that FTD arises from disruptions in semantic processing, whereas the dysexecutive hypothesis attributes FTD to deficits in executive control. Both hypotheses are supported by converging evidence from behavioral and neuroimaging studies^9–15^. Importantly, positive FTD highlights the intrinsic coupling between language and thought. Disruption in semantic processing or associative mechanisms can rise to derailment and tangentiality reflecting excessive or aberrant conceptual connections. These findings suggest that semantic and syntactic deficits, alongside executive dysfunction, constitute key cognitive underpinnings of positive FTD. Meta-analyses have indicated that positive FTD is significantly associated not only with semantic processing and syntactic comprehension deficit, but also with impairments in executive functioning, including working memory, planning, and inhibitory control^16,17^. Neuroimaging studies have identified abnormal activation in regions essential for coherent speech production, which play a central role in the manifestation of positive FTD in schizophrenia^18,19^. Furthermore, resting-state functional connectivity analyses have revealed altered interactions between temporal seed regions implicated in positive FTD and cortical areas involved in executive and higher-order cognitive processes, including working memory and social cognition^15^. Collectively, these findings suggest that positive FTD reflects an interplay between dyssemantic and dysexecutive dysfunction.

Negative FTD, in contrast to positive FTD, is characterized by slowed thinking and impoverished speech. However, this impoverishment reflects not only verbal output deficits but reduced intrinsic thought generation, leading to difficulty mobilizing concepts and associations necessary for coherent language organization. This pattern suggests that while thought processes are impaired, core language system remains relatively intact, supporting the dissociation between language and thought^20^. Slowed thinking, linked to diminished higher-order functions such as executive control, is therefore consistent with the dysexecutive hypothesis. Moreover, the weak correlation between positive and negative FTD scores supports distinct neurocognitive impairments^16,21,22^. Structural and functional neuroimaging studies link negative FTD to deficits in regions integral to the executive control network, including gray matter reductions in the medial frontal cortex, dorsolateral and orbitofrontal cortices, and anterior cingulate cortex^23^, consistent with dysexecutive mechanisms.

In this study, we examined resting-state functional connectivity alterations within the language and executive control networks and their associations with positive and negative FTD in schizophrenia. We employed a novel dynamic “meta-networking” framework derived from meta-network theory^24^, which emphasizes spatiotemporal network properties during rest^25^, providing insights into intrinsic network dynamics and functional segregation.. To minimize confounding effects from chronic disease, comorbidities, and long-term environmental influences, which can significantly affect both fMRI and clinical outcomes, we focused on first-episode schizophrenia patients recruited from two independent imaging centers. To further assess the clinical translational value of these imaging metrics, we included a third independent imaging center, where individuals with first-episode schizophrenia underwent a two-week standardized medication course, and MRI data were acquired once clinical stabilization was achieved.

## Result

### Demographics and clinical information

We conducted, a multicenter dynamic meta-networking investigation to delineate alterations in the language and executive control networks and their associations with positive and negative FTD. Resting-state fMRI data were pooled from three independent cohorts: a discovery cohort comprising150 patients, 175 healthy controls; a validation cohort including183 patients and 109 healthy controls; and a short-term medication cohort of 71 patients rescanned after two weeks of standardized treatment, each with matched controls.

Across all sites, there were no significant differences in age and sex distribution between HCs and individuals with schizophrenia for the three sites (Table 1). Both the discovery and validation cohorts consisted of drug-naive patients, resulting in relatively elevated PANSS scores. Notably, patients in the discovery cohort demonstrated more severe clinical profile, including higher positive FTD, negative FTD, and PANSS without FTD scores. In contrast, patients in the short-term medication cohort exhibited significant symptom improvement following treatment, with the lowest PANSS, positive FTD, and negative FTD scores across the three cohorts.

**Table 1.**
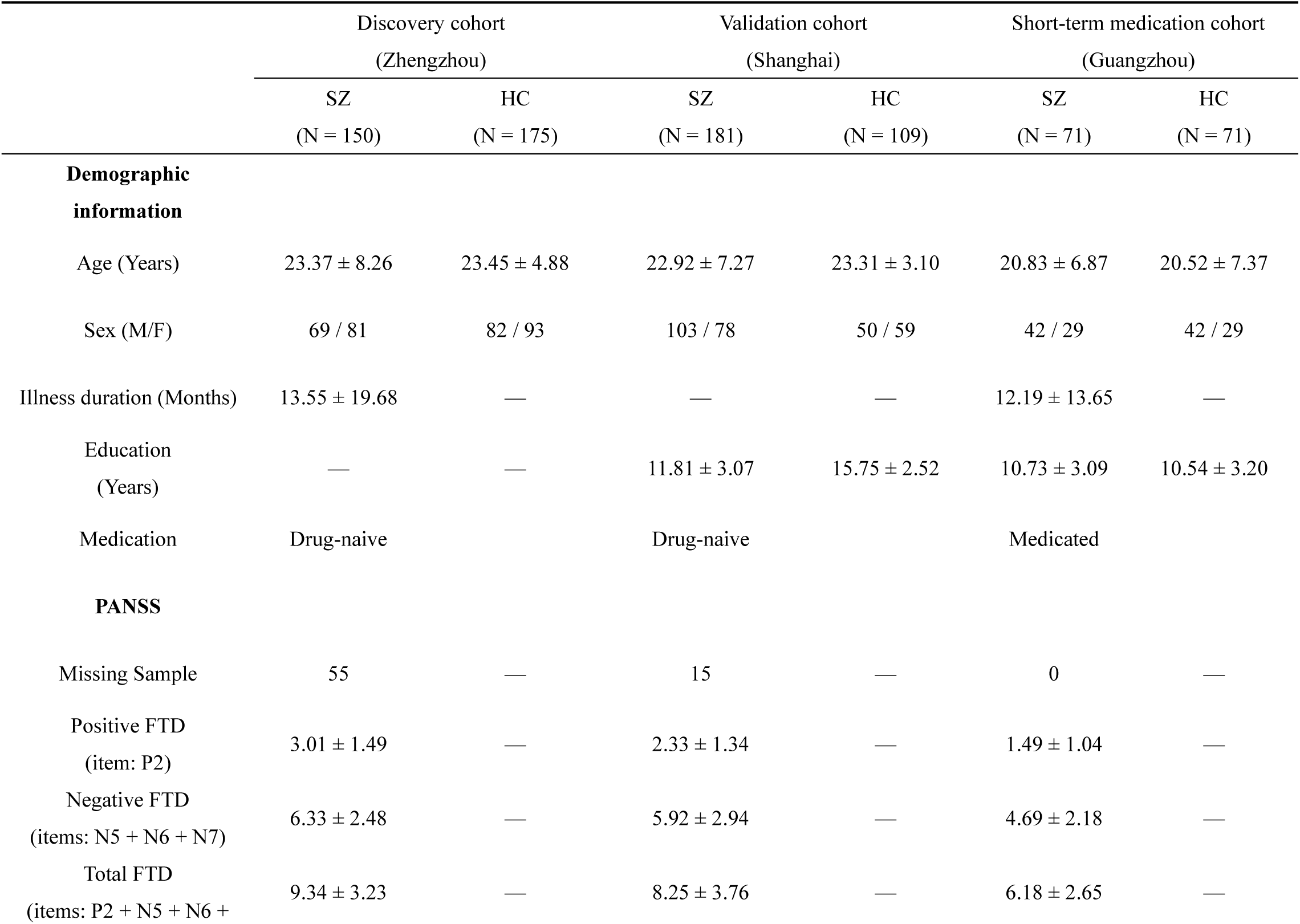
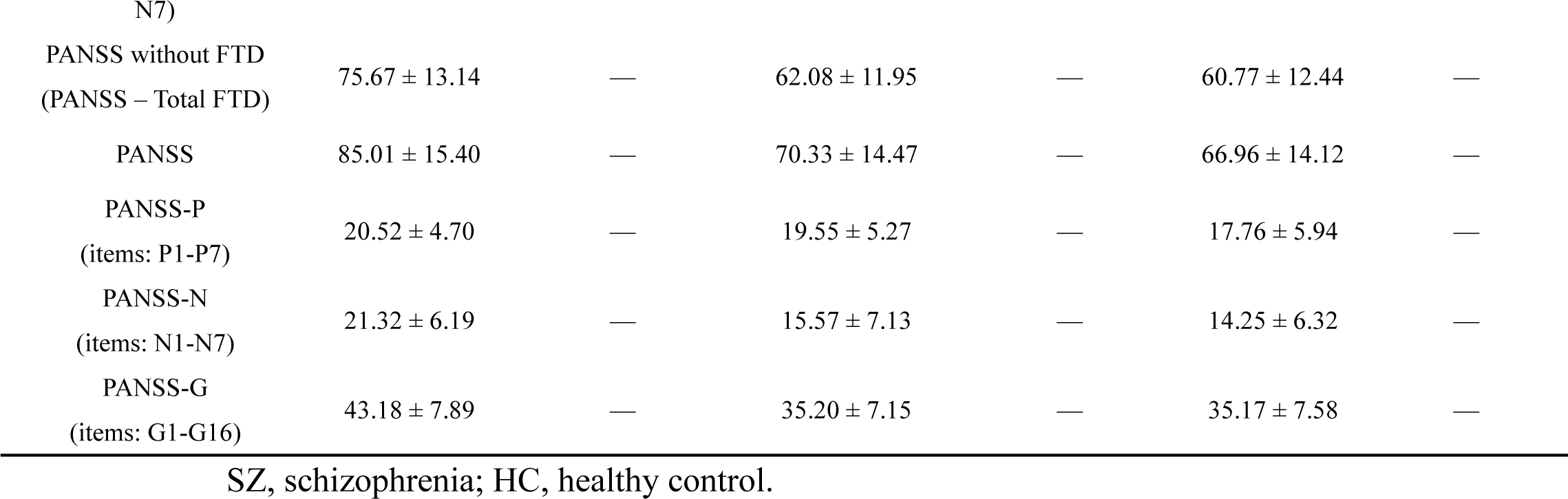
Demographic and clinical information.

### Correlations between positive and negative FTD

We examined the distribution of positive and negative FTD across the three cohorts. In the discovery cohort, positive FTD displayed a moderate distribution, ranging from minimal to pronounced symptom severity, whereas negative FTD was consistently elevated with substantial inter-individual variability. The validation cohort reproduces this pattern: positive FTD was skewed toward milder presentations yet extended into more severe range, while negative FTD remained broadly elevated. Crucially, in the short-term medication cohort, both symptom domains shifted toward minimal severity and displayed markedly reduced dispersion, reflecting substantial clinical improvement after two weeks of treatment.

Notably, across all cohorts, positive and negative FTD scores were positively correlated, with weak-to-moderate effect sizes (discovery cohort: *r* = 0.278, *p* = 0.006; validation cohort: *r* = 0.467, *p* < 0.001; short-term medication cohort: *r* = 0.264, *p* = 0.026) (Fig. 1).

**Figure 1.**
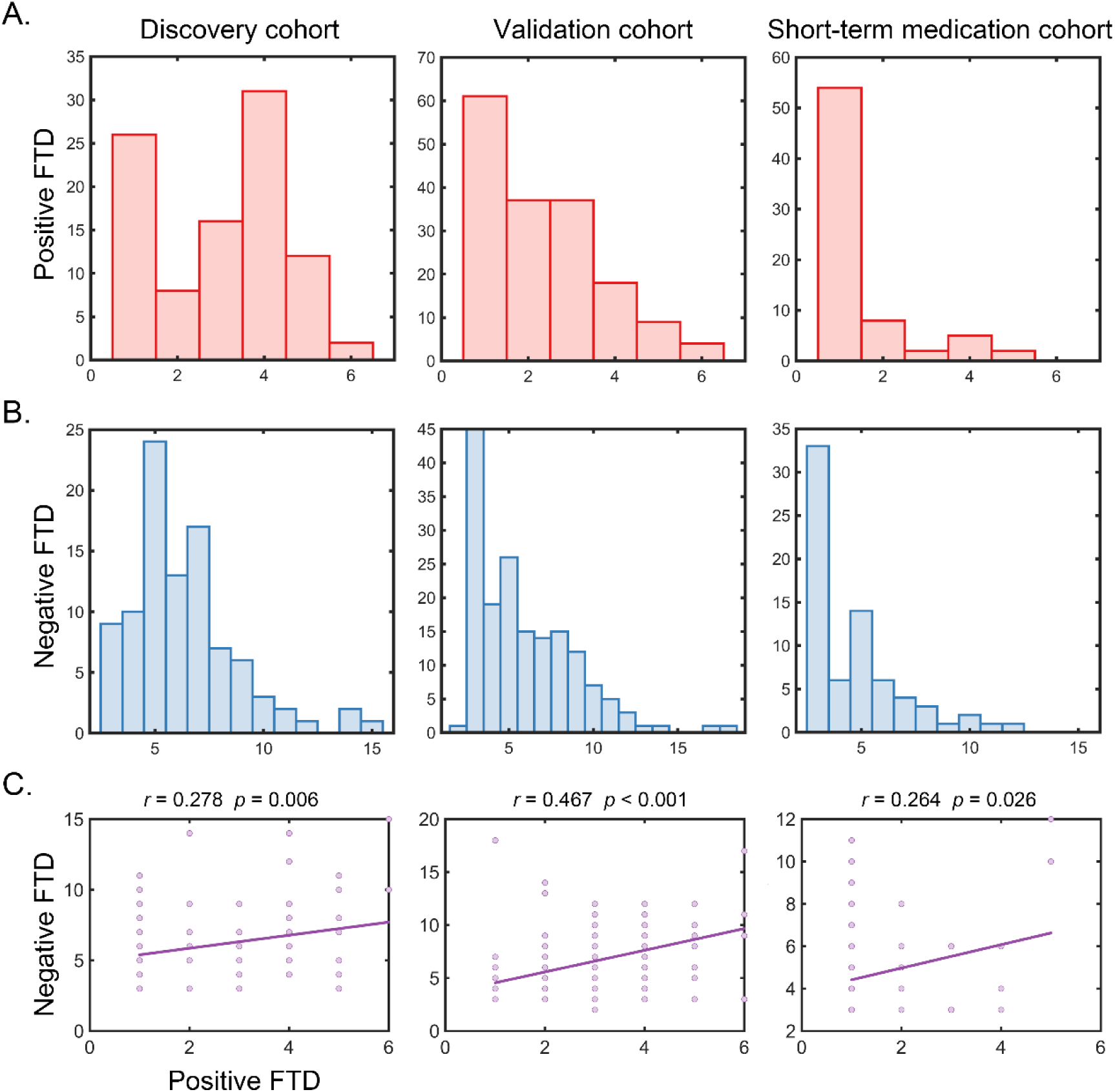
Distribution of positive and negative FTD scores and their correlations. Participants were recruited from three imaging sites: the discovery cohort (Zhengzhou), the validation cohort (Shanghai), and the short-term treatment cohort (Guangzhou). (A) Distribution of positive FTD scores. (B) Distribution of Negative FTD scores. (C) Scatterplots depicting correlations between positive FTD and negative FTD.

### Meta-networking states of the language and executive control networks

A univariate generalized autoregressive conditional heteroskedasticity (GARCH)-based dynamic conditional correlation (DCC) model was applied to estimate frame-wise, time-varying connectivity matrices, which were subsequently partitioned into recurring meta-states using k-means clustering (Fig. 2).

**Figure 2.**
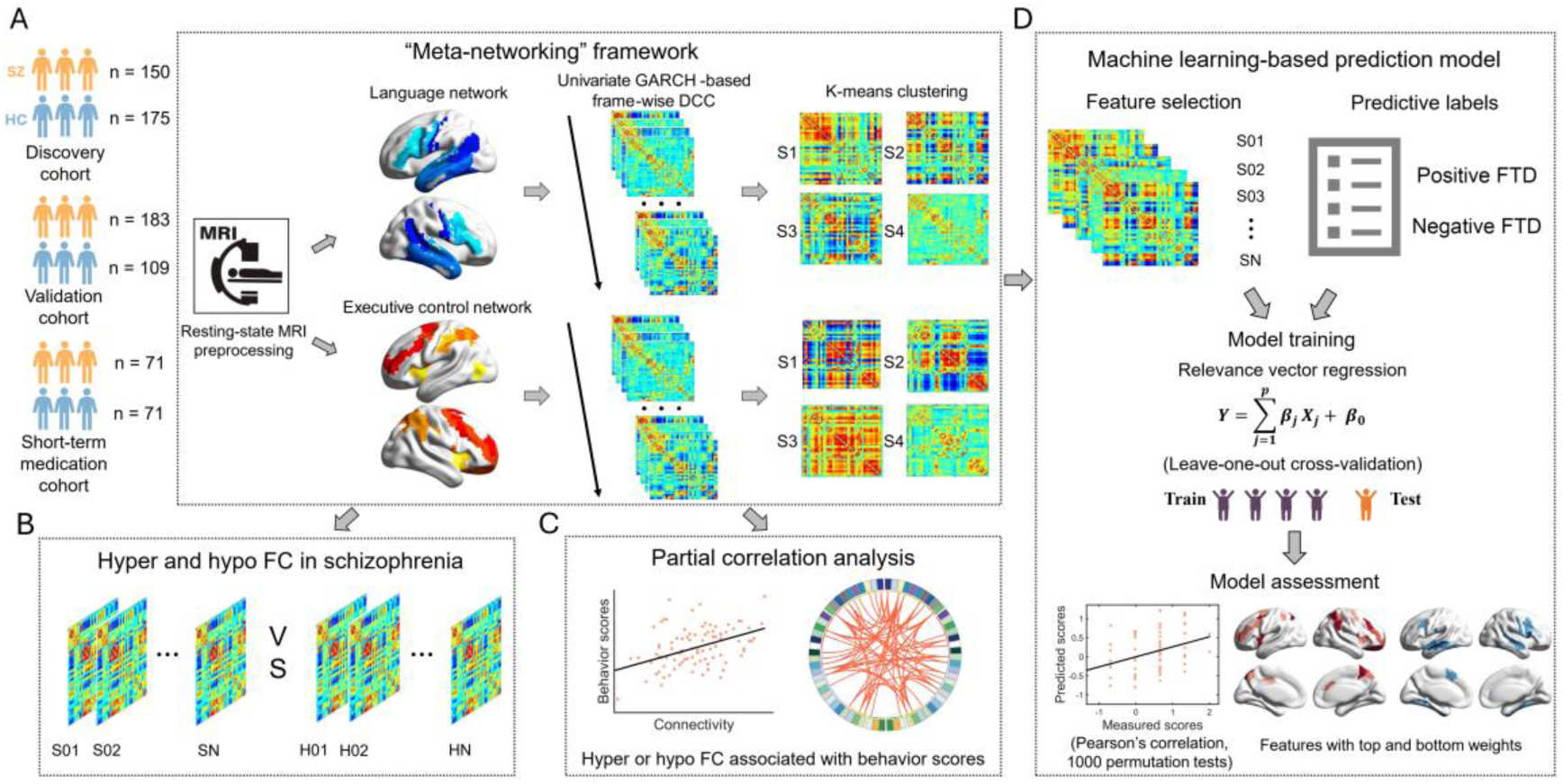
Flowchart of the “Meta-Networking” Framework. (A) Resting-state fMRI data were acquired from three independent imaging sites, including both HC and individuals with SZ. After preprocessing, dynamic conditional correlation (DCC) and k-means clustering were applied to derive recurrent the meta-states of the language and executive control networks (B) Two-sample t-tests were conducted for each identified state to assess patterns of hyper- and hypo-connectivity in SZ relative to HC. (C) Partial correlation analysis was performed to identify state-specific connections associated with positive or negative FTD. (D) Predictive models for positive and negative FTD were constructed using relevance vector regression (RVR) with a leave-one-subject-out cross-validation framework. Model significance was evaluated with 1,000 permutation tests, and for significant models (p < 0.05), the top 5% of most informative features were extracted. dFC: dynamic functional connectivity; DCC: dynamic conditional correlation; GARCH: Generalized Autoregressive Conditional Heteroskedasticity.

Consistent with prior findings^25^, four meta-networking states with distinct functional connectivity patterns were identified within the language network in both HCs and individuals with SZ (Fig. 3). Across all four states, nodal strength conformed to exponentially truncated power-law distributions, indicating the presence of a subset of nodes with disproportionately high connectivity, commonly referred to as hubs. The top 30 nodes by nodal strength within each state were designated as hubs. State-dependent hub distributions were observed in the first three states. In State 1, hub nodes were primarily located in the posterior sections of the STG and STS, as well as the ventral sections of the PrG and PoG. State 2 was characterized by hubs predominantly distributed in the IFG, ventral PrG, posterior STG, and FuG. State 3 featured hubs within the MTG, IPL, and IFG. In contrast, State 4 represented a weakly connected state in which no hubs were identified.

**Figure 3.**
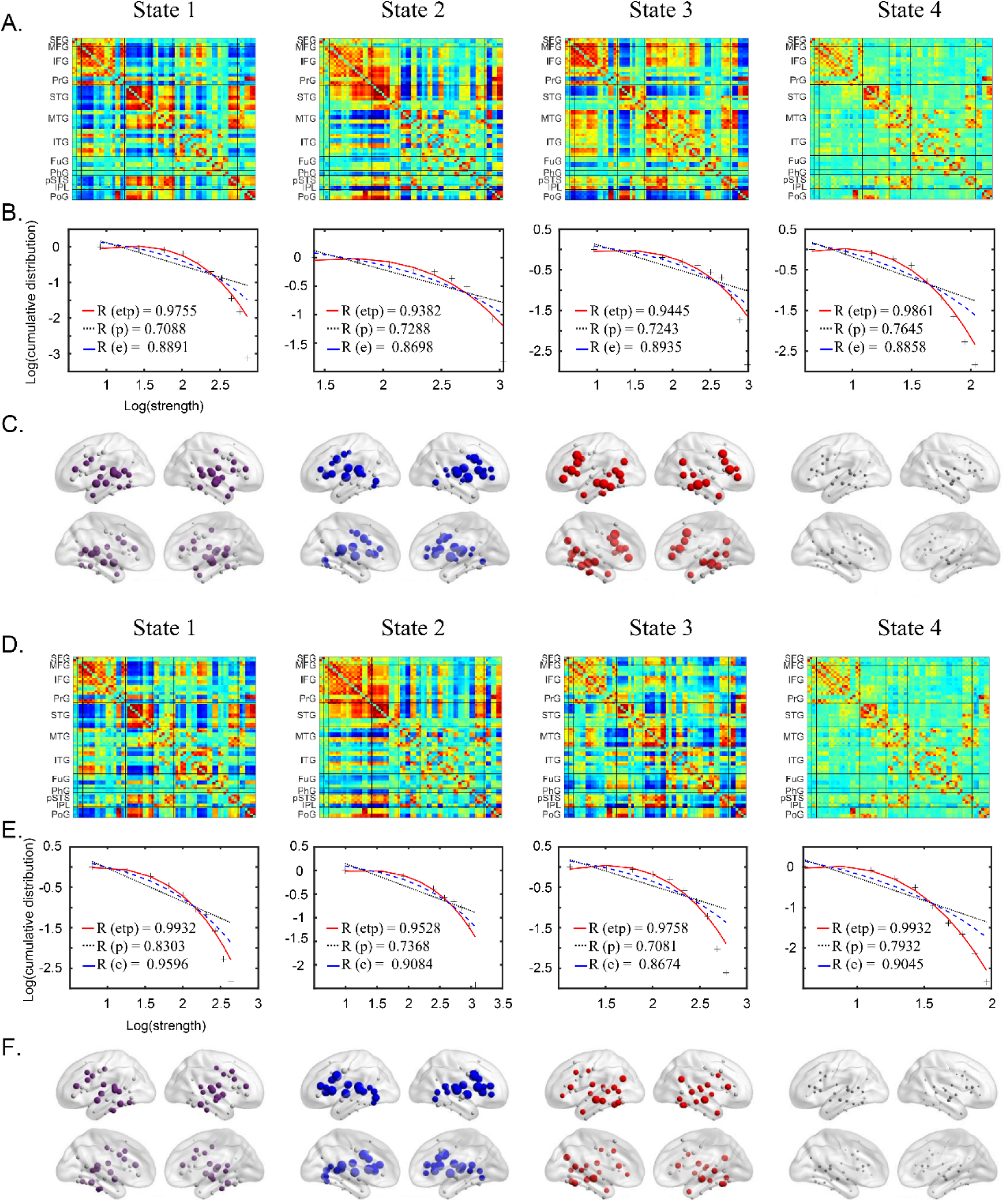
The four meta-networking states of the language network. **(A & D)** Connectivity matrices of the four states in HCs and SZ from the discovery cohort. Four temporally reoccurring states with distinct functional connectivity patterns were identified. **(B & E)** Log-log plots of the cumulative nodal strength distributions. The plus sign (black) represents observed data, the solid line (red) is the fit of the exponentially truncated power-law distribution, 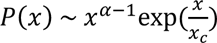, the dashed line (blue) is an exponential distribution, 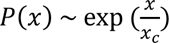, and the dotted line (black) is a power-law, *P*(*x*) ∼ *x*^α−1^. *R*^2^ was calculated to assess the goodness of fit, with higher values indicating better model fitting. The exponentially truncated power-law is the best fit for all four states. **(C & F)** nodal strength distributions. The first 30 nodes with the highest nodal strength were defined as hubs.

**Figure 4.**
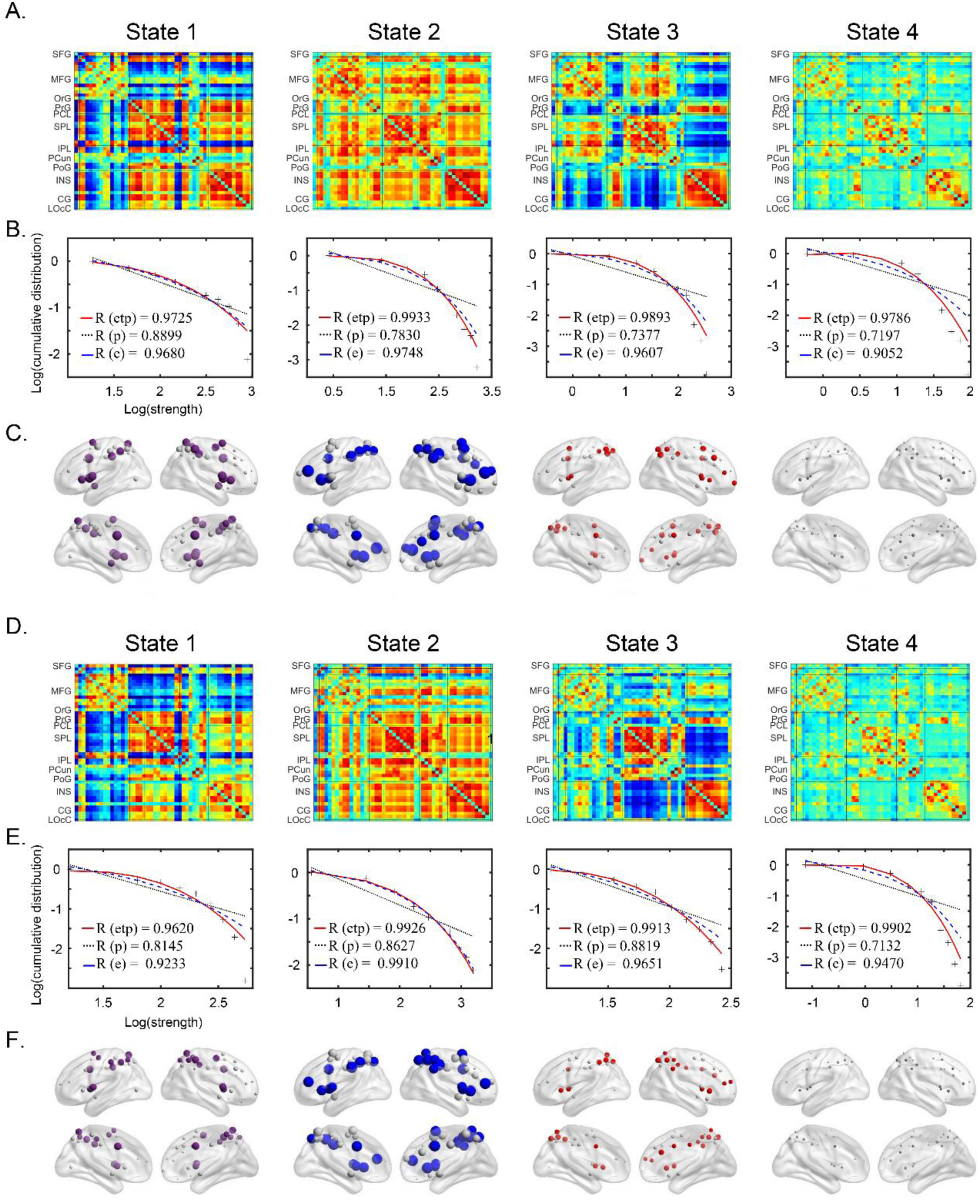
The four meta-networking states of the executive control network. **(A & D)** connectivity matrices of the four states in HCs and schizophrenia from the discovery cohort. Four temporally reoccurring states with distinct functional connectivity patterns were identified. **(B & E)** Log-log plots of the cumulative nodal strength distributions. An exponentially truncated power-law is the best fitting for all four states. **(C& F)** nodal strength distributions. The first 20 nodes with the highest nodal strength were defined hubs.

To evaluate the functional significance of state-dependent hub distributions, we conducted term-based meta-analytic activation analyses. Five functional terms were analyzed: speech perception, inner speech, phonological processing, speech production, and semantic processing. spatial correspondence between the state-specific hub distribution and the term’s meta-analytic activation pattern was weak to moderate, with Dice coefficients ranging from 0.21-0.48 in HCs, and 0.16-0.46 in individuals with SZ (Supplementary Fig. 2). These findings indicate that language network exhibits a domain-separated organization of resting-state dynamics. State 1 was most likely to be implicated in speech perception and phonological processing, State 2 with speech production processing, and State 3 with semantic processing.

Within executive control network, four meta-networking states were identified in both HCs and individuals with SZ. Across all states, nodal strength followed exponentially truncated power-law distributions. In State 1, hubs (defined as the top 20 nodes ranked by nodal strength) were localized primarily to the bilateral CG, PrG, and bilateral SPL (Fig 5). In State 2 was characterized by hubs were predominantly found in the bilateral IPL, SPL, right MFG, and insula. In State 3, hubs were distributed across the bilateral SPL, insula, and CG. in contrast, State 4 represented a weakly connected state, in which no hub was identified.

**Figure 5.**
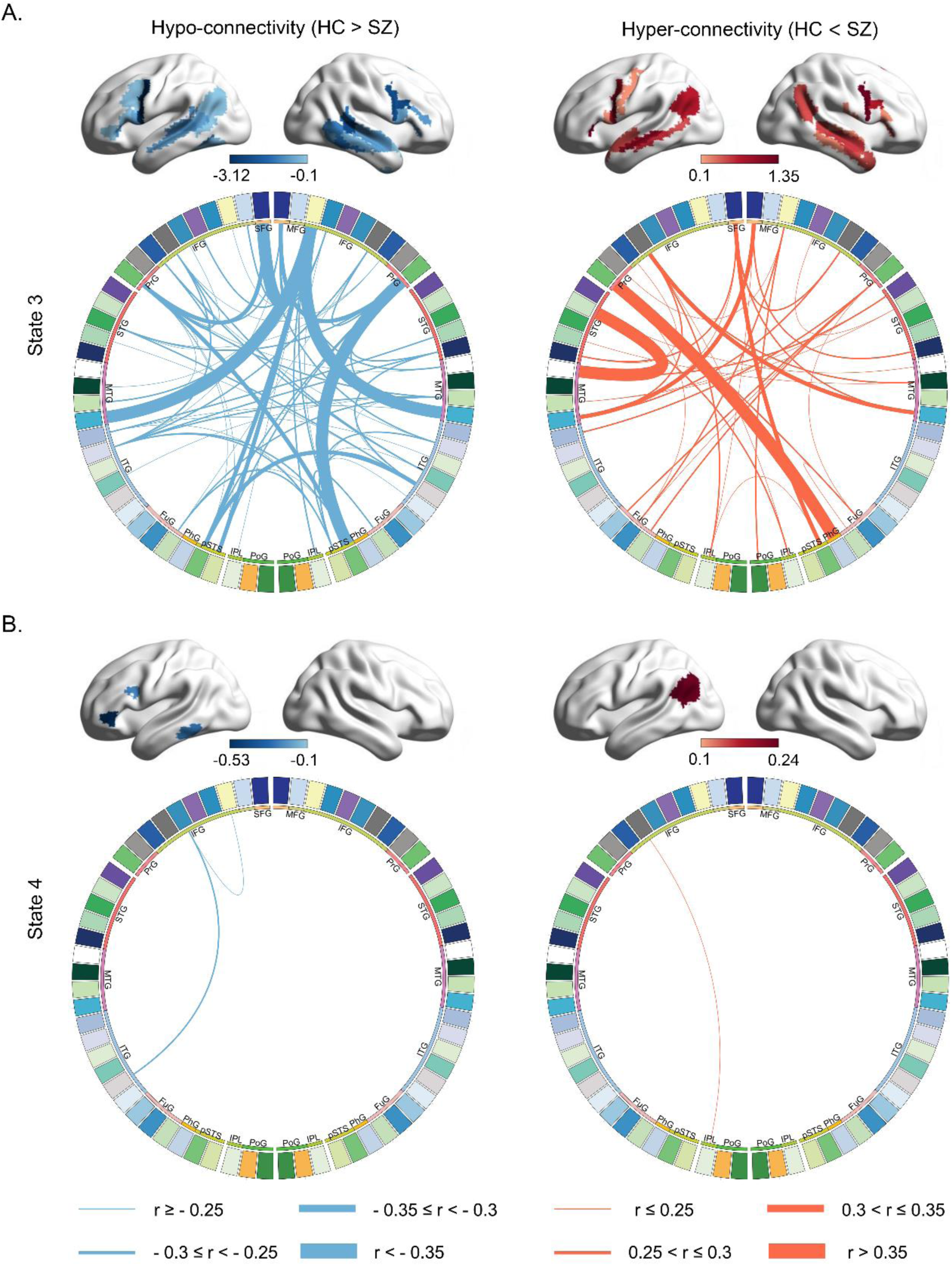
Hypo- and hyper-connectivity of the language network and their associations with positive FTD. (A) Circo plots and cortical surface renderings of state 3, depicting NBS-significant hypo- and hyper-connectivity linked to positive FTD (B) Circo plots and cortical renderings of state 4, showing significant associations between connectivity alterations and positive FTD severity.

Term-based meta-analytic activation analyses were also performed for the executive control network. Three functional terms were assessed: dorsal attention network, ventral attention network, frontoparietal control network. Network parcels were defined using the Brainnetome Atlas. Weak to moderate correspondence was again observed between to the three networks were defined from state-specific hub distribution and the term’s activation pattern, with Dice coefficients of 0.07-0.31 in HCs, and 0.05-0.41 in individuals with SZ (Supplementary Fig. 3). State 1, hubs overlapped primarily with the dorsal and ventral attention networks, State 2 hubs overlapped mainly with the dorsal attention and frontoparietal control networks, whereas State 3 hubs showed a comparable overlap across all three networks.

### Prediction of positive and negative FTDs using meta-networking states

Using a relevance vector regression (RVR) framework with leave-one-out cross-validation (LOOCV), we tested whether meta-networking connectivity patterns could predict (individual differences) FTD severity.

In the language network, predictive model based on the combined connectivity patterns across all four states significantly predicted positive FTD scores (*r* = 0.487, *p* < 0.001). When models were constructed separately for each state, the semantic-related State 3 (*r* = 0.401, *p* = 0.011) and the baseline state (*r* = 0.342, *p* = 0.015) significantly predicted positive FTD scores (Table 2; Supplementary Fig. 4).

**Table 2.** Machine learning-based prediction model accuracies and significance.

|  |  |  | Positive FTD |  | Negative FTD |  |
| --- | --- | --- | --- | --- | --- | --- |
|  |  |  | *r* | *p* | *r* | *p* |
| Discovery cohort | Language network | State 1 | 0.269 | 0.12 | 0.205 | 0.122 |
|  |  | State 2 | 0.197 | 0.073 | 0.162 | 0.112 |
|  |  | State 3 | **0.401** | **0.011** | -0.137 | 0.789 |
|  |  | State 4 | **0.342** | **0.015** | -0.011 | 0.508 |
|  |  | State 1+2+3+4 | **0.487** | **< 0.001** | 0.141 | 0.225 |
|  | Executive control network | State 1 | 0.187 | 0.1 | **0.326** | **0.004** |
|  |  | State 2 | **0.351** | **0.043** | 0.211 | 0.123 |
|  |  | State 3 | 0.144 | 0.136 | **0.31** | **0.013** |
|  |  | State 4 | **0.319** | **0.041** | 0.174 | 0.21 |
|  |  | State 1+2+3+4 | 0.217 | 0.15 | **0.34** | **0.024** |

In the executive control network, the combined connectivity patterns across all four states significantly predicted negative FTD scores (r = 0.34, p = 0.024). State-specific analyses further revealed significant prediction of negative FTD by State 1 (*r* = 0.326, *p* = 0.004) and State 3 (*r* = 0.31, *p* = 0.013). in addition, State 2 (*r* = 0.351, *p* = 0.043) and State 4 (*r* = 0.319, *p* = 0.041) significantly predicted positive FTD severity (Table 2; Supplementary Fig. 5).

### Prediction of positive and negative FTDs using meta-states derived from a combined language–executive control network parcellation

Meta-state features were derived from a combined language-executive control parcellation, predictive models did not achieve significance for either positive FTD (*r* = 0.119, *p* > 0.05) or negative FTD (*r* = 0.027, *p* > 0.05).

### Hypo- and hyper-connectivity associated with the positive FTD

For both network, case-control comparisons were performed using two-sample t-tests with network-based statistics (NBS) correction (Edge *p* = 0.01, Component *p* = 0.01).

In the language network, both hypo- and hyper-connectivity were observed across all four meta-states relative to HC (Supplementary Fig. 6). Given that the semantic-processing (State 3) and baseline (State 4) meta-states significantly predicted positive FTD scores, we next examined brain–behavior associations. NBS-significant links were extracted and tested using partial Pearson correlations (controlling sex, age, head motion with 10000 permutations). State 3 exhibited the strongest disruptions: a subset of hypo-connected links correlated negatively with positive FTD severity whereas a subset of hyper-connected links correlated positively. In State 4, greater positive FTD severity was associated with reduced connectivity in the IFG and ITG, and increased connectivity in the IPL (Fig. 5; Supplementary Fig. 7).

In the executive control network, both hypo- and hyper-connectivity were also observed across all four meta-states (Supplementary Fig. 8). Connectivity alterations in meta-states 2 and 4 were predominantly associated with positive FTD. Specifically, reduced connectivity in the MFG, IPL, and insula correlated with more severe positive FTD, whereas increased connectivity in the CG, SPL, and PrG correlated with milder positive FTD symptoms. (Fig. 6; Supplementary Fig. 9).

**Figure 6.**
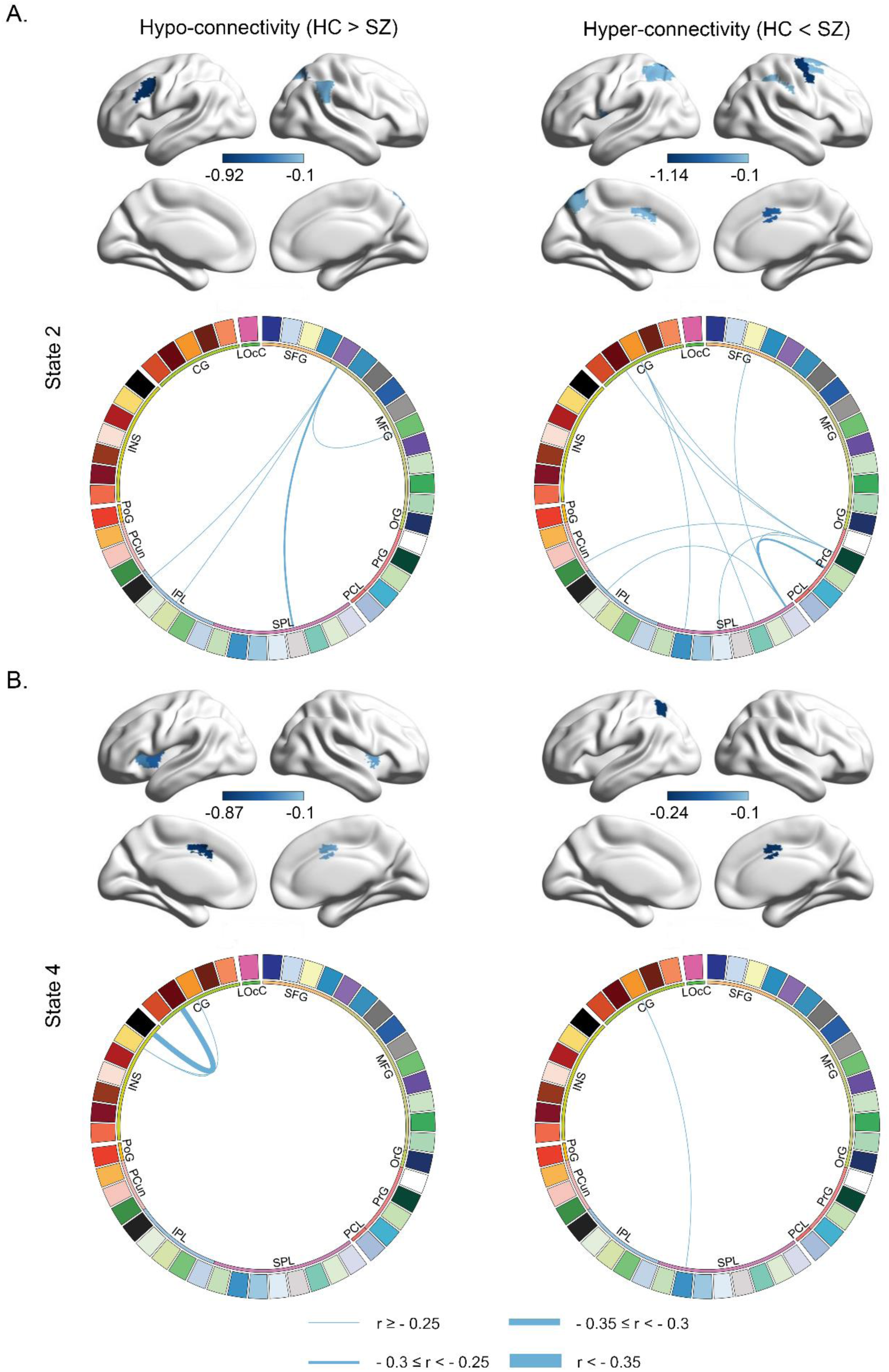
Hypo- and hyper-connectivity of the executive control network and their associations with positive FTD. (A) Circo plots and cortical renderings of state 2, depicting NBS-significant hypo- and hyper-connectivity associated with positive FTD. (B) Circo plots and cortical renderings of state4, showing connectivity changes significantly associated with positive FTD severity.

### Hypo- and hyper-connectivity associated with the negative FTD

In States 1 of the executive control network, hypo-connectivity among the MFG, insula, CG, and right PrG was negatively correlated with negative FTD, whereas hyper-connectivity between the MFG, CG, IPL and PCL was positively correlated with negative FTD (Fig 7; Supplementary Fig 14). In State 3, hypo-connectivity involving the insula was negatively correlated with negative FTD, while hyper-connectivity between the MFG and CG was positively correlated with negative FTD (Fig 7; Supplementary Fig 10).

**Figure 7.**
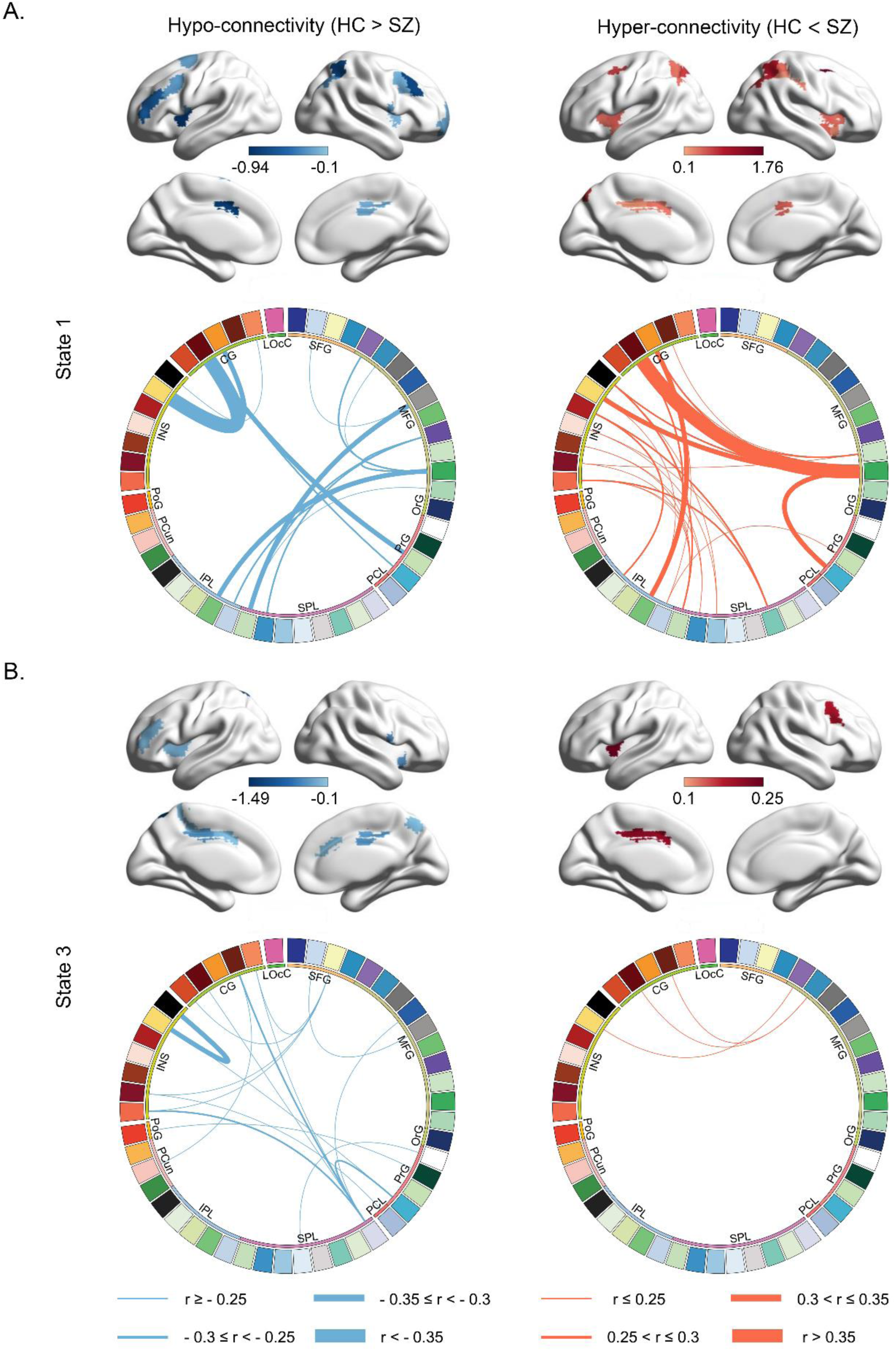
Hypo- and hyper-connectivity of the executive control network and their associations with negative FTD. (A) Circo plots and cortical surface renderings of state1 showing NBS-significant hypo- and hyper-connectivity associated with negative FTD. (B) Circo plots and cortical surface renderings of state 3 depicting significant association with negative FTD.

### Static FC analysis

To assess whether static connectivity captured similar effects, we applied an analogous analysis pipeline using static FC (Supplementary Fig. 23 - 25, Supplementary Table 6). Predictive analyses revealed that only static FC within the language network significantly predicted positive FTD scores (*r* = 0.36, *p* = 0.004). No significant predictive effect was observed for the executive control network on either positive (*r* = 0.197, *p* > 0.05) or negative FTD (*r* = 0.114, *p* > 0.05).

### Results of the Validation dataset

Consistent with the discovery dataset, both hypo- and hyper-connectivity were observed in SZ across the language (Supplementary Fig. 2, 6 & 11) and executive control networks (Supplementary Fig. 3, 8 & 12). Predictive models using features from all states (r = 0.436, p < 0.001) and from each state individually (State 1: *r* = 0.261, *p* = 0.009; State 2: *r* = 0.477, *p* < 0.001; State 3: *r* = 0.3, *p* = 0.022; State 4: *r* = 0.421, *p* = 0.004) significantly predicted positive FTD (Supplementary Table 3) significantly predicted positive FTD. Within States 3 and 4, significant negative correlations were observed between hypo-connectivity in the temporoparietal junction, STG, MTG, and PoG, and positive FTD (Supplementary Fig. 13).

For the executive control network, hypo-connectivity in MFG and IFG was significantly negatively correlated with negative FTD (Supplementary Fig. 14). Hypo-connectivity in MFG and hyper-connectivity in parietal lobe were positively associated with positive FTD (Supplementary Fig. 15). Models using features from all states (*r* = 0.331, *p* = 0.009) and from state 1 (*r* = 0.272, *p* = 0.046) significantly predicted negative FTD (Table 2). Moreover, atypical connectivity features in State 2 predicted positive FTD (*r* = 0.355, *p* = 0.007) (Supplementary Table 3). collectively, these resutls support the stability of our findings across cohorts indicated that our results were stable across different cohorts.

### Normalization of FTD-related hypo- and hyper-connectivity after two-weeks medication

Similar patterns of hypo- and hyper-connectivity associated with positive and negative FTD were observed after two weeks of standardized medication (Supplementary Fig 16 - 20). The predictive model based on connectivity features from all states of the executive control network significantly predicted negative FTD scores (*r* = 0.395, *p* = 0.004), underscoring the robustness of these features in capturing negative FTD. In contrast, language network connectivity did not significantly predict positive FTD after treatment (*r* = 0.036, *p* > 0.05) (Supplementary Table 3).

Building on the identification of shared network disruptions, we further investigated whether these abnormalities were attenuated following short-term antipsychotic treatment. We identified the top 10 connections in the discovery cohort that exhibited the highest correlations with either positive or negative FTD, representing the most significant differences in connectivity associated with FTD. To assess functional connectivity normalization, we calculated the effect sizes for these connections at both the discovery and short-term medication cohorts, comparing patients with HCs. The analysis revealed generally larger effect sizes at the discovery cohort than at the short-term medication cohort across both the language and executive control networks (Fig. 8), indicating greater deviation from normal connectivity prior to treatment. Specifically, at the discovery cohort, language network effect sizes spanned 0.381-0.934 (mean ± s.d. = 0.587 ± 0.173) in hypo-connected regions and 0.312-0.788 (mean ± s.d. = 0.430 ± 0.141) in hyper-connected regions. At the short-term medication cohort, they spanned 0.291-0.708 (mean ± s.d. = 0.478 ± 0.153) in hypo-connected regions and 0-0.337 (mean ± s.d. = 0.127 ± 0.101) in hyper-connected regions. Likewise, in the executive control network, effect sizes in hypo-connected regions at the discovery cohort spanned 0.368–3.279 (mean ± s.d. = 0.839 ± 0.875) versus 0–0.743 (mean ± s.d. = 0.148 ± 0.244) at the short-term medication cohort, and in hyper-connected regions 0.421–2.501 (mean ± s.d. = 0.862 ± 0.657) at the discovery cohort versus 0–0.188 (mean ± s.d. = 0.033 ± 0.062) at the short-term medication cohort. Among those edges, treatment-related reductions in effect size ranged from −0.319–3.231. This reduction in effect sizes after two weeks of medication suggests a general trend toward normalization of the FTD-related functional abnormalities, particularly in the connections previously identified as most disrupted across sites.

**Figure 8.**
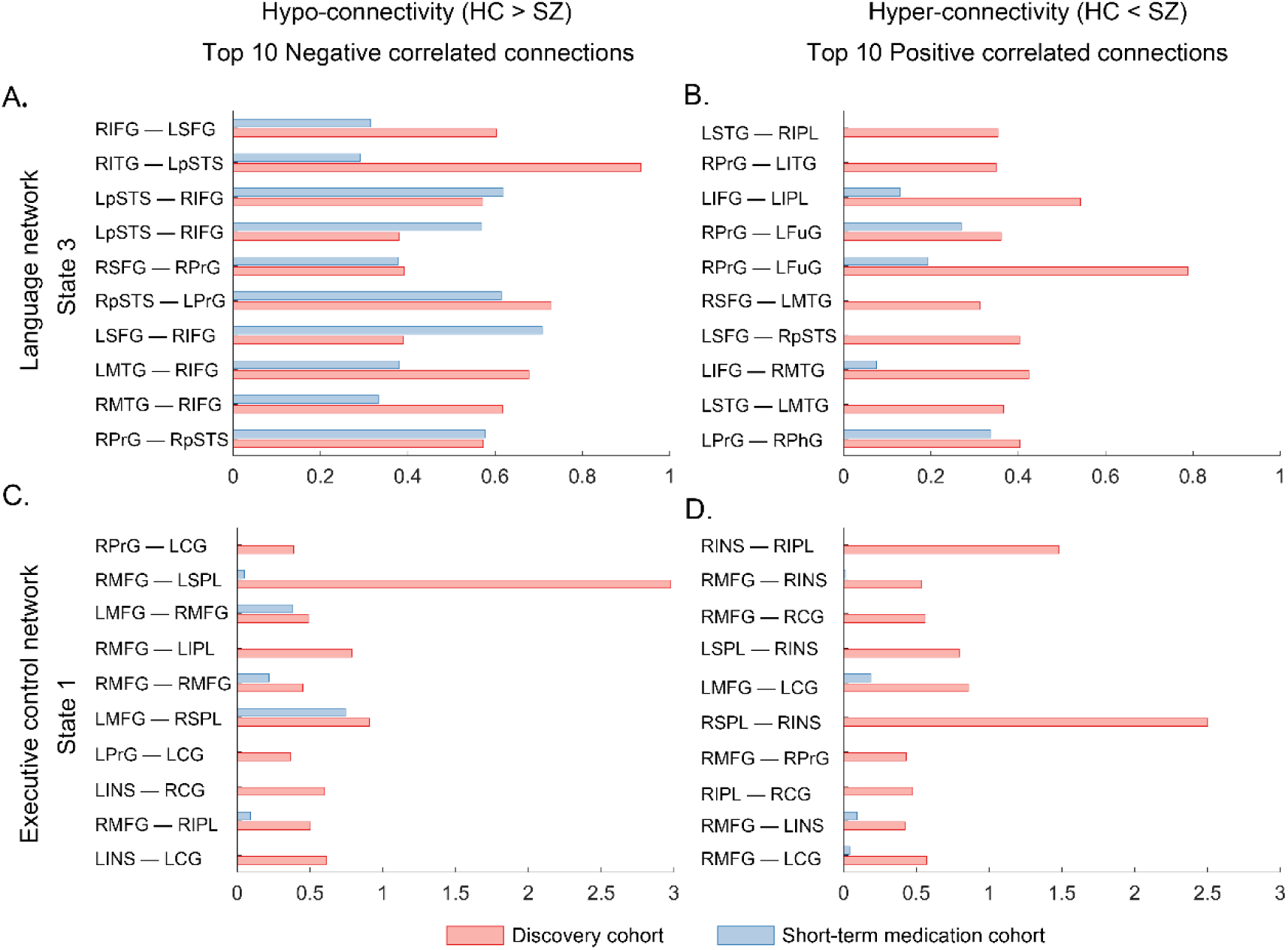
Short-term medication effects on FTD-associated brain connectivity. (A & B) Effect sizes for hypo- and hyper-connectivity in state 3 of the language network associated with positive FTD. (C & D) Effect sizes for hypo- and hyper-connectivity regions in state 1 of the executive control network associated with negative FTD.

### Shared hypo- and hyper-connectivity between sites

To determine whether network disruptions extended beyond FTD-specific effects, we identified, for each meta-state, the connectivity alterations common to discovery and the short-term medication cohorts.

In language network, consistent disruptions were observed across the IFG, PrG, FuG, and temporal lobe. State-specific network disruptions included: State 1, the IFG, MTG, ITG, and pSTS; State 2, the FuG, IFG, ITG, and pSTS; State 3, the PrG, ITG, IFG, and FuG; State 4, the STG and PrG (Supplementary Fig. 21).

In the executive control network, shared disruptions were observed in: State 1, (right IPL, insula, CG, PrG, and the left SFG, MFG, SPL, and PoG); State 2, (right insula, MFG, SFG, CG, and left PoG); State 3, (IPL, PrG, SPL, MFG); and State 4, (bilateral insula and CG) (Supplementary Fig. 22).

These consistent network interruptions were also significantly correlated with non-FTD PANSS, implicating these connections in broader SZ symptomatology. In the language network, significant correlations with non-FTD PANSS were found in: State 1(SFG, ITG and MTG); State 2 (MFG, pSTS); State 3 (MFG, IFG, FuG and ITG), State 4 (STG) (Supplementary Table 4). In the executive control network, significant correlations included: state1(SPL and CG); State 2(insula and SPL); State 3 (CG, MFG and SPL); State 4 (CG, MFG and SPL) (Supplementary Table 5).

## Discussion

In this study, we provide evidence for a polarity-dependent model of the network mechanisms underlying positive and negative FTD in SZ. Using a meta-networking framework, we demonstrated that positive FTD is associated with altered connectivity across specific meta-states of both the language and executive control networks, providing convergent support for the dyssemantic and dysexecutive hypotheses. In contrast, negative FTD was confined to disruption in two meta-states of the executive control network, consistent with the dysexecutive hypothesis. Of particular translational relevance, these network alterations normalized following short-term treatment, underscoring their promise as dynamic biomarkers with potential clinical utility.

Clinically, positive FTD manifest as language disturbance including derailment and tangentiality, whereas negative FTD is reflected in diminished informational content of speech^26^. Consistent with the discriminant validity of these dimensions, we observed a weak correlation between positive and negative FTD, replicating prior findings^27,28^. This weak correlation persisted even after brief pharmacological intervention, reinforcing the conceptual independence of the two symptom clusters. Prior work has similarly documented distinct association of positive and negative FTD with cognitive processes, and diagnostic specificity.^29–31^. Recent studies^13,32^, including the present study, further delineates the distinct neurobiological mechanisms underlying these two forms of FTD.

Our meta-networking analyses identified four distinct states of the language network during resting state, consistent with prior studies^25,33^. Both univariate correlation and model-based prediction analyses converged on the finding that only State 3, associated with semantic processing, was significantly linked to positive FTD. In hypo-connectivity patterns, regions most negatively associated with positive FTD included the bilateral pSTS, MTG, left PrG, and right IFG. A previous study has delineated two positive FTD dimensions (“disorganization” and” incoherence”) and one negative dimension (’emptiness’)^34^, supported by the identification of two white matter subnetworks corresponding to these factors. Critically, the disorganization subnetwork showed substantial overlap with cortical regions involved in language and speech production, underscoring its pivotal role in language dysfunction^35^. In line with this, the left MTG and pSTS have been identified as core hubs of the disorganization subnetwork implicated in schizophrenia-related FTD^35^. Moreover, the severity of positive FTD has been inversely correlated with activity in the IFG, left STG, and MTG^36–38^, regions fundamental to coherent speech production, auditory-speech processing, and semantic integration^39,40^. Conversely, hyper-connectivity associated with positive FTD was observed in the bilateral MTG, SFG, right PhG, left FuG, and PrG. Prior research has similarly reported increased neural activity in the FuG among individuals with positive FTD, as well as positive correlations between positive FTD and (activation of the?) PhG, anterior FuG, and PrG^32,37,41–43^. These regions are consistently implicated in semantic feature processing^12,44,45^. Collectively, these findings suggest dysfunction within the language network as a central pathophysiological mechanism underlying positive FTD.

Similarly, our analyses identified four distinct states of the executive control network during the resting-state. State 1 was characterized by both hypo- and hyper-connectivity patterns. Across univariate correlation and multivariate predictive modeling, State 1 emerged as selectively associated with negative FTD in both the discovery and validation datasets. Within this state, hypo-connectivity was negatively correlated with negative FTD in the bilateral CG, MFG, IPL, and left INS. Structural and functional abnormalities in the insula have previously been linked to a higher severity of negative FTD^46–48^. Moreover, task-based fMRI studies have reported deactivation in the MFG and CG that inversely correlated with FTD severity^49–52^. These regions are critically engaged in cognitively demanding operations, including response conflict monitoring^53^, cognitive control engagement^54^, top-down inhibitory modulation^55^, mnemonic competition detection, and retrieval-induced forgetting^56^. Disruptions of these functions may contribute (directly) to impairments in thought organization and speech coherence. Concurrently, hyper-connectivity within State 1 showed positive associations with negative FTD in the bilateral CG, INS, SPL, right MFG, and IPL. Task-based fMRI studies have highlighted the SPL’s role in attentional switching^57^,with negative FTD demonstrating positive correlations with SPL activation under high switching demands^58^. Although the SPL has been less frequently implicated in FTD, our findings provide novel evidence for its functional dysregulation in the pathophysiology of negative FTD.

Additionally, machine-learning analyses revealed that State 2 of the executive control network significantly predicts positive FTD, suggesting that executive control network abnormalities also contribute to the pathogenesis of positive FTD. Prior seed-based functional connectivity studies have linked positive FTD to higher-order cognitive processes^15^, with structural abnormalities observed in the cortical regions of the executive control network^59,60^. Our meta-networking findings converge with this evidence, highlighting. the close interplay between semantic impairments and executive dysfunction in positive FTD. The integration of fundamental language processes with higher-order executive functions appears essential for generating coherent and contextually comprehensive speech.

Our results highlight polarity-dependent neural substrates of FTD symptomatology and underscore a fundamental dissociation between language and thought. The language network primarily encompasses key frontal regions such as the MFG and IFG, as well as temporal regions such as the STG, MTG, and STS^61–64^. This network supports the storage of abstract linguistic knowledge and the execution of computations required for accessing word meanings and implementing combinatorial syntactic and semantic processes^65^. By contrast, thought is mediated by the multiple-demand network, comprising bilateral frontal and parietal cortices along with medial prefrontal and posterolateral inferior temporal domain-general areas^66,67^. These regions are constantly engaged in a broad range of high-level cognitive operations, including executive control, novel problem-solving, and logical reasoning^68–71^. Importantly, convergent evidence indicates that complex thought processes are not dependent on language^20^. Functional mapping methods demonstrate that language-specific regions exhibit minimal activation during tasks such as mental arithmetic, working-memory maintenance, or cognitive control^72–74^. Lesion studies provide further support: individuals with global aphasia, despite profound deficits in language comprehension and production, often retain the capacity to perform mathematical calculations and solve logical problems^75,76^. Conversely, individuals with SZ may show impairments in abstract thinking and reasoning despite relatively preserved linguistic functions^77^. Together, these findings suggest that language is neither necessary nor sufficient for thought, reinforcing the conceptual dissociation between linguistic processing and higher-order cognition.

### Conclusion

Using a meta-networking framework for dynamic network analysis, we demonstrate that positive FTD is characterized by altered connectivity across specific meta-states of both the language and executive control networks. By contrast, negative FTD is linked exclusively to dysfunction within two executive control network meta-state, consistent with the dysexecutive hypothesis. Notably, these polarity-dependent network disruptions normalized following short-term anti-psychotic treatment and coincided with clinical improvement in FTD symptoms.

## Methods

### Participants

The first-episode schizophrenia was diagnosed using the Diagnostic and Statistical Manual of Mental Disorders, Fourth Edition criteria and the MINI. Participants were recruited from three independent imaging sites to ensure the robustness and generalizability of the findings. The discovery cohort, recruited from the First Affiliated Hospital of Zhengzhou University, included 150 first-episode, drug-naïve individuals diagnosed with schizophrenia and 175 HCs. An independent replication cohort, recruited from Shanghai Mental Health Center, comprised 183 first-episode, drug-naïve individuals with schizophrenia and 109 HCs. To investigate whether neuroimaging biomarkers associated with Positive or Negative FTD normalize after clinical treatment, a third cohort, the Affiliated Brain Hospital of Guangzhou Medical University, was included. This cohort, consisting of inpatients who had received two weeks of antipsychotic medication, comprised 71 individuals with first-episode schizophrenia and 71 HCs.

Exclusion criteria included: 1) past or current neurological diseases or mental disorders due to physical diseases; 2) obvious active somatic diseases; and 3) MRI contraindications including the presence of metallic implants, braces, or claustrophobia.

The HCs were recruited through local advertisement postings that had no any lifetime psychiatric or neurological diagnoses and had no family history of any psychiatric disorders.

Data were collected from the National Key R&D Program of the Ministry of Science and Technology of China between 2016 and 2021. The research ethics committees of the above-mentioned three hospitals approved the study. Written informed consent was obtained from all participants during recruitment. For participants younger than 18 years, informed consent was provided by both the participants and their parents. All procedures conducted in this study complied with the ethical standards of the relevant national and institutional committees on human experimentation and the 1975 Declaration of Helsinki, revised in 2008.

### Psychopathology Assessment and FTD assessment

All patients completed the Chinese versions of the Positive and Negative Symptoms Scale (PANSS)^78^, which consists of 30 items divided into 3 psychopathology subscales: positive symptoms (PANSS-P; items P1-P7), negative symptoms (PANSS-N; N1-N7), and general psychopathological symptoms (PANSS-G; G1-G16). Structured clinical interviews were conducted by senior psychiatrists who completed the training required for this type of investigation. The interrater reliability of the PANSS, as per the ratings of the trained interviewers, ranged from 0.76 to 0.92.

To quantify the severity of FTD symptoms, we derived symptom scores based on subscales from PANSS. Four distinct scores were computed for each patient: total FTD, positive FTD, negative FTD, and total PANSS excluding FTD. The assessment of FTD symptoms was based on four specific PANSS items: P2, N5, N6, and N7. Among these, P2 (conceptual disorganization) was used to measure positive FTD, while negative FTD was quantified as the sum of N5 (difficulty in abstract thinking), N6 (lack of spontaneity and flow of conversation), and N7 (stereotyped thinking).

### MRI data acquisition

MRI was scanned for all subjects within 3 days before or after clinical assessment, Resting-state functional MRI data in the discovery cohort were acquired on a GE 3.0 T MRI system. Whole-brain coverage was achieved using a gradient-echo single-shot echo-planar imaging (EPI) sequence with the following parameters: repetition time (TR) = 2000 ms, echo time (TE) = 30 ms, flip angle (FA) = 90°, field of view (FOV) = 220 × 220 mm², acquisition matrix = 64 × 64, slice thickness = 4.0 mm, inter-slice gap = 0.5 mm, number of slices = 32, and number of volumes = 180. In addition, T1 structural data were collected in the sagittal plane with a spoiled gradient-echo sequence (3D T1WI) using the following parameters: TR / TE = 8.2 / 3.2 ms; FA = 12°; FOV = 256 × 256 mm²; matrix size = 256 × 256; slice thickness = 1.0 mm; inter-slice gap = 0 mm; number of slices = 188.

Individual MRI data in short-term medication cohort were collected on a Philips Achieva 3.0 T TX MRI system (Philips Healthcare, Netherlands) equipped with an eight-channel head coil array. Specifically, the rs-fMRI data were acquired axially using a fast field echo-echo planar imaging sequence. The parameters were as follows: 33 slices, TR = 2000 ms, TE = 30 ms, FOV = 220 × 220 mm², FA = 80°, matrix = 64 × 64, slice thickness = 4 mm, number of volumes = 240, voxel size = 3.44 × 3.44 × 4 mm^3^, and inter-slice gap = 0.6 mm. High resolution T1-weighted images were collected with a sagittal T1-weighted 3D turbo field echo sequence: TR = 820 ms, TE = 380 ms, FOV = 256 × 256 mm², voxel size = 1 × 1 × 1 mm^3^, matrix = 256 × 256, 188 sagittal slices, slice thickness = 1 mm Imaging data in the validation cohort were acquired with 3.0 T Siemens Verio MRI scanner (Siemens AG, Erlangen, Germany) with a 32-channel head coil located. The parameters were as follows: 37 slices, TR = 2500 ms, TE = 30 ms, FOV = 100 × 100 mm², FA = 90°, matrix = 64 × 64, slice thickness = 3.5 mm. To reduce movement, a foam padding was placed around the subject’s head. T1 structural data were acquired using a 3D T1-weighted magnetization prepared rapid gradient echo (MPRAGE) sequence (TR = 2530 ms, TE = 2.56 ms, FOV = 256 × 256 mm², voxel size = 1 × 1 × 1 mm^3^, FA = 7°, matrix = 256 × 256, 192 sagittal slices, slice thickness = 1 mm).

### Processing of MRI Data

The functional data were preprocessed using the following steps: (1) Removal of the first 10 volumes to allow for signal stabilization; (2) slice timing correction to account for differences in acquisition time across slices; (3) head motion correction, ensuring that motion did not exceed 3 mm or 3°); (4) coregister of functional images to the corresponding anatomical T1 images; (5) Segmentation and normalization of T1 images into MNI space; (6) Normalization of functional images to the MNI space; (7) Spatial smoothing with a Gaussian kernel (full-width-at-half-maximum = 6 mm); (8) Linear detrending to remove low-frequency drifts; (9) Regression of nuisance signals using 36 parameters (36 parameters, including the x, y, and z translations, and rotations + WM/CSF/global time courses [9 parameters], plus their temporal derivatives [9 parameters] and the quadratic terms of 18 parameters)^79,80^; and (10) temporal band-pass filtering (0.01-0.1 Hz).

### Network definition of language and executive control

The putative network parcels for the two networks were defined using the Human Brainnetome Atlas^81^. This atlas comprises 246 regions, parcellated based on white matter structural connectivity patterns. To identify functional relevance for each parcel, Fan and colleagues conducted meta-analyses utilizing behavioral domain and paradigm-class meta-data labels from the BrainMap database. From these analyses, we extracted all parcels that exhibited significant activation associated with language-related or executive control-related behavioral domains or paradigm classes.

For the language network, 33 cortical parcels in the left hemisphere and 15 in the right hemisphere were selected. These regions closely correspond to those identified by other researchers, reflecting consistency in delineating the language network. Given the symmetric connectivity patterns observed during the resting state and the involvement of both hemispheres in language processing, 19 homologs in the right hemisphere and one homologous parcel in the left hemisphere were additionally included. This approach resulted in a symmetric language network atlas (Supplementary Fig. 1) comprising 68 cortical parcels, including the superior, middle, and inferior frontal gyrus (SFG, MFG, and IFG, respectively); the ventral parts of the precentral gyrus (PrG) and postcentral gyrus (PoG); the superior, middle and inferior temporal gyrus (STG, MTG, and ITG, respectively); the posterior superior temporal sulcus (pSTS); the inferior parietal lobule (IPL); the fusiform gyrus (FuG); and the parahippocampus gyrus (PhG).

For the executive control network, we selected parcels corresponding to the dorsal attention network, ventral attention network, and frontal-parietal network. To ensure independence from the language network, parcels overlapping with language network nodes were excluded. As a result, the executive control network (Supplementary Figure 1) was defined with 50 parcels, encompassing regions such as the superior and middle frontal gyrus (SFG and MFG), orbital gyrus (OrG), precentral gyrus (PrG) and paracentral lobule (PCL). Additionally, it included the superior and inferior parietal lobule (SPL and IPL), precuneus (PCun), postcentral gyrus (PoG), insula cortex (INS), cingulate gyrus (CG), and lateral occipital cortex (LOcC).

For ROIs, such as MFG (label: IFJ_L, ID: 17), that exhibit both language and execution control functions, network assignment is based on the meta-analysis maximum.

The coordinates (in Montreal Neurologic Institute space) and meta results of each parcel for the two networks are summarized in Supplementary Tables 1 and 2.

### Meta-network construction using DCC

The dynamic conditional correlation (DCC) approach was utilized to construct framewise time-varying networks^82^. As a variant of the multivariate generalized autoregressive conditional heteroscedasticity (GARCH) model, DCC estimates dynamic correlations between multiple time series by incorporating these time-varying variances. All DCC model parameters are estimated using quasi-maximum likelihood methods, eliminating the need for ad hoc parameter tuning. Briefly, The DCC model operates in two sequential steps. First, each parcel’s time series is standardized via a univariate GARCH (1,1) model to remove temporal heteroscedasticity. Then, an exponentially weighted moving average (EWMA) is applied to those standardized residuals to produce a time-varying correlation matrix.

### Identification of Meta-Networking States Using K-Means Clustering

To identify meta-network states (i.e., temporally recurring states), the K-means clustering algorithm was employed to partition the dynamic functional connectivity matrices into a set of non-overlapping clusters. After determining the number of clusters (*k*), each dFC matrix was assigned to a specific cluster (state) based on the similarity of its spatial pattern to the cluster centroid. The clustering process aimed to minimize the sum of distances between individual dFC matrices and the centroid of their respective clusters, with the centroid serving as the representative pattern for that cluster.

The optimal number of clusters (*k*) was determined using the elbow criterion, which evaluates the ratio of within-cluster distances to between-cluster distances^83^. To quantify the distance between points and centroids, the L1 distance function (Manhattan distance) was applied. Finally, each dFC matrix from every time frame was assigned to one of the identified clusters, representing a specific meta-networking state.

### Nodal strength distribution and hubs

To investigate the presence of hubs in each state, we calculated the nodal strength distribution *P(k)* (Supplementary Methods).

### Functional relevance of state-dependent hub distributions

To investigate the functional relevance of each meta-state, we performed term-based meta-analyses. For the language network, fMRI-based activations of language- and speech-related terms from the Neuroquery platform, including speech perception, inner speech, phonological processing, speech production, and semantic processing, were queried. For the executive control network, terms were derived from the three clusters of spatially distinct executive function networks identified in prior research: the dorsal attention network, the ventral attention network, and frontoparietal control network^84^.

The functional relevance of hubs was evaluated using Dice coefficients, which quantify the spatial overlap between the binary images of hub distributions and meta-analytic results. The Dice coefficient is defined as: Dice = 2∗ (hub ∗ meta)/(hub + meta). Considering the lateralized nature of meta-analytic activations, Dice coefficients were calculated specifically for the whole brain, only in left hemisphere, and only in the right hemisphere.

### Machine learning-based state prediction model

To examine the relationship between reorganized dFC states and FTD, we developed a relevance vector regression (RVR) model^85,86^. Prediction accuracy was evaluated using leave-one-out cross-validation (LOOCV), which calculates the Pearson correlation coefficient between predicted and actual labels (Supplementary Methods). Model significance was assessed through 1000 permutation tests.

### Statistical analysis

Two-sample t-tests were performed to analyze age, FTD scores, PANSS scores as well as nodal and global topological metrics. Sex differences were assessed using Pearson’s chi-square test.

Hypo- and hyper-connectivity were evaluated using two-sample t-tests, with statistical significance corrected through network-based statistics (NBS) at an edge-level threshold of *p* = 0.01 and a component-level threshold of *p* = 0.01. Nodal topological properties were corrected for multiple comparisons using the False Discovery Rate (FDR), with a corrected significance threshold of *p* = 0.05. Cohen’s d values, representing the effect size of the group comparisons, were computed from the t-statistic.

For the single state exhibiting significant prediction, the NBS was also used to identify brain-behavior associations that were evaluated by calculating partial Pearson correlation coefficients. The significance of correlation was assessed through 10000 permutation tests. Confounding factors, sex, age, and head motion were controlled.

## Complementary analysis

As with dynamic FC, we applied an analogous analysis pipeline to static FC to compare the sensitivity of both methods.

## Data/code availability statement

The datasets analyzed are upon request to the correspondence authors.

All codes used to process structural images and calculate network measures are from publicly available toolboxes (DPABI^87^, https://github.com/Chaogan-Yan/DPABI;

NaDyNet^88^, https://github.com/yuanbinke/Naturalistic-Dynamic-Network-Toolbox)

## Conflict of Interest Statement

No competing financial interests exist.

## Acknowledgements

The study is supported by the National Social Science Foundation of China (Grant No. 20&ZD296), the Key-Area Research and Development Program of Guangdong Province (Grant No. 2019B030335001), the National Natural Science Foundation of China (Grant Nos. 32400862, 82151314, and 82472092), the Research Center for Brain Cognition and Human Development, Guangdong, China (Grant No. 2024B0303390003), the Clinical Research Plan of Shanghai Hospital Development Center (Grant No. SHDC12022113), grants from Guangzhou Science and Technology Bureau, Municipal (enterprise) and Institute (Grant Nos. 2024A03J0298, 2024A03J0219), the STI 2030—Major Projects (Grant No. 2021ZD0200500), Guangzhou Research-oriented Hospital.

## Credit authorship contribution statement

Z Hu, JS Chen: formal analysis, writing – original draft, funding acquisition, visualization; BK Yuan, XC Tang, JJ Wang, LP Cao: supervision, project administration, writing – original draft.

Other authors: writing – original draft.

## Notes

### Competing Interest Statement

The authors have declared no competing interest.

