## Supplementary Figure 1, Supplementary Figure 2, Supplementary Figure 3, Supplementary Figure 4, Supplementary Figure 5, Supplementary Figure 6 for "Dynamic Meta-Networking Identifies Distinct Network Correlates of Positive and Negative Formal Thought Disorder in Schizophrenia"

#### Supplementary Methods

##### Nodal strength and distribution

The nodal strength for a given node was calculated by summing up the correlation coefficients of all the suprathreshold connections that connected with this node. To determine whether there were hubs in each state, we calculated the nodal strength distribution $P\ (k)$. Three possible forms of distribution were fitted to the probability of nodal strength: a power-law, $P\ (x) \sim x^{\alpha-1}$; and exponential, $P\ (x) \sim \exp\ ({}^{x}/_{x_c})$; and an exponentially truncated power-law, $P\ (x) \sim x^{\alpha-1} \exp\ ({}^{x}/_{x_c})$. Among these, the exponentially truncated power-law distribution is commonly observed in human brain networks. This distribution characterizes long-tailed, broad-scale topologies, where a substantial proportion of network connectivity is concentrated on a subset of nodes, referred to as hubs[5–7]. Given the distinct connectivity patterns and strengths across the dynamic functional connectivity (dFC) states, hubs within each state were identified as nodes with strengths exceeding the mean value of the respective state's nodal strength distribution.

**Machine learning-based state prediction model**

We used a relevance vector regression (RVR) model to estimate FTD severity based on the recognized dFC states. RVR is a Bayesian framework for learning sparse regression models. In RVR, only some samples (smaller than the training sample size), termed the "relevance vectors," are used to fit the model:

$$y(x) = \sum_{i=1}^{m} \omega_i \tau_i + \epsilon,$$

where $\tau_i$ are basis functions, and $\epsilon$ is normally distributed with mean 0 and variance β. RVR uses training data to build a regression model:

$$y = \theta\omega + \epsilon,$$

where $y = [y_1, y_2, \cdots y_n]^T$, $\theta = [\theta_1, \theta_2, \cdots \theta_n]$, $\theta_i = [\tau_i(x_1), \tau_i(x_2), \cdots \tau_i(x_n)]^T$. Each vector $\theta_i$, consisting of the values of the basis function τi for the input vectors, is a relevance vector.

The model parameters $\beta$ were found by using the maximum likelihood estimates from the conditional distribution: $p\ (y \mid \alpha, \beta) = N\ (y \mid 0, C)$ ,where the $C = \beta I_n + \Phi A^{-1} \Phi^T$. To make the RVM favor sparse regression models, prior distributions were assumed for both $\omega_i$ and $\beta^{-1}$, i.e. $p\ (w_i \mid \alpha_i) = N\ (0, \alpha_i^{-1})$.The ways of treating priors, however, lead to the same relevance vector machine construction.

The leave-one-out cross-validation (LOOCV) was used to estimate prediction accuracy. During each iteration of LOOCV, one patient was designated as the test sample, while the remaining patients were used to train the prediction model. The predicted score for the test sample was then derived using its feature matrix.

Significance was assessed through 1000 permutation tests. For each permutation, the prediction labels (e.g., the positive or negative FTD scores) were randomly shuffled, and the RVR prediction process was repeated. This resulted in a distribution of random accuracies. The P-value was calculated as: P = (number of permutation tests < actual accuracy + 1) / (number of permutation tests + 1). The features for each prediction model included the subject's median matrix for a specific state or their combination.

### Supplementary Figures

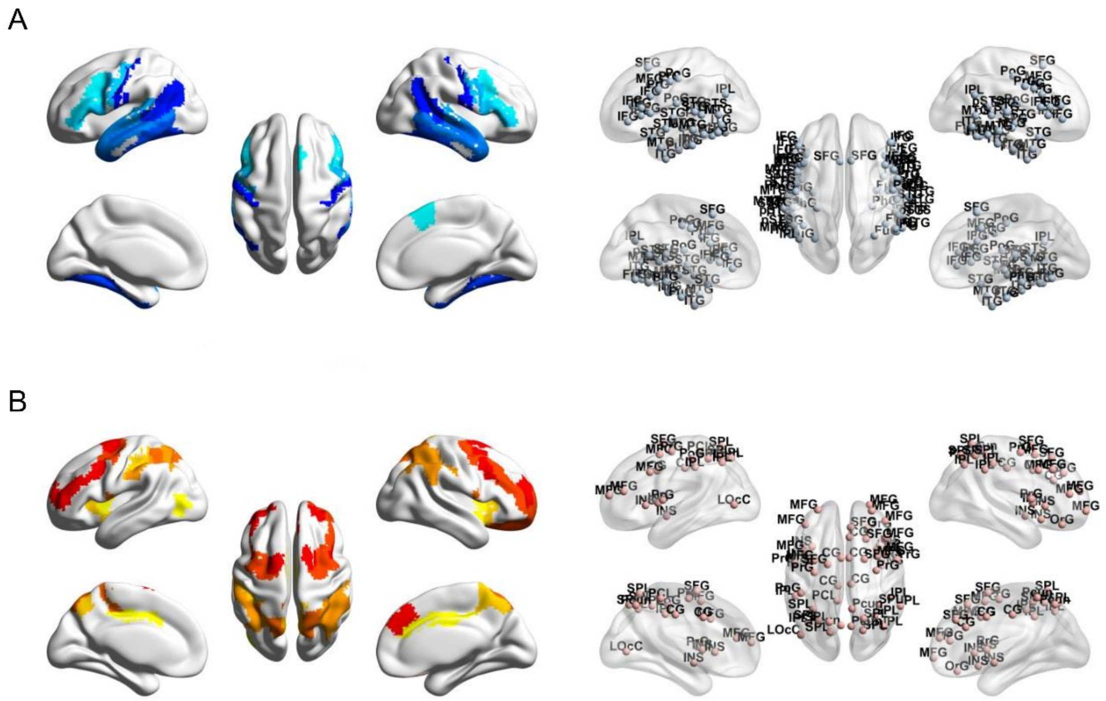


**Supplementary Figure 1.** (A). The language network consisting of 68 regions. (B). The executive control network consisting of 50 regions. The two networks were defined based on the Brainnetome Atlas (n = 246, https://atlas.brainnetome.org/bnatlas.html) according to each node's meta-analytic behaviorial domains and paradigm classes.

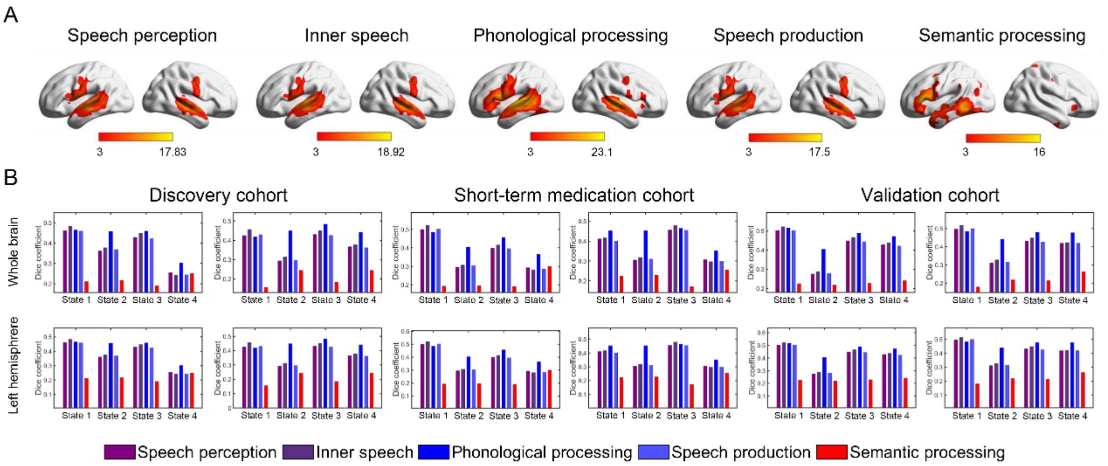


**Supplementary Figure 2. Functional relevance of hubs in the first three states of language network.** (A). Meta-analysis results for speech perception, inner speech, phonological processing, speech production, and semantic processing obtained from ‘NeuroQuery’ (*https://neuroquery.org/*) (Dockes et al., 2020). Each map was thresholded at $Z = 3$ (a typical value used by NeuroQuery) for illustrative purposes and only positive results were displayed. (B). Dice coefficients between the binary images of network nodes and meta-analysis results. Given the left-lateralized activations in the meta-analysis, Dice coefficients were calculated both at the whole-brain level and with in the left hemisphere.

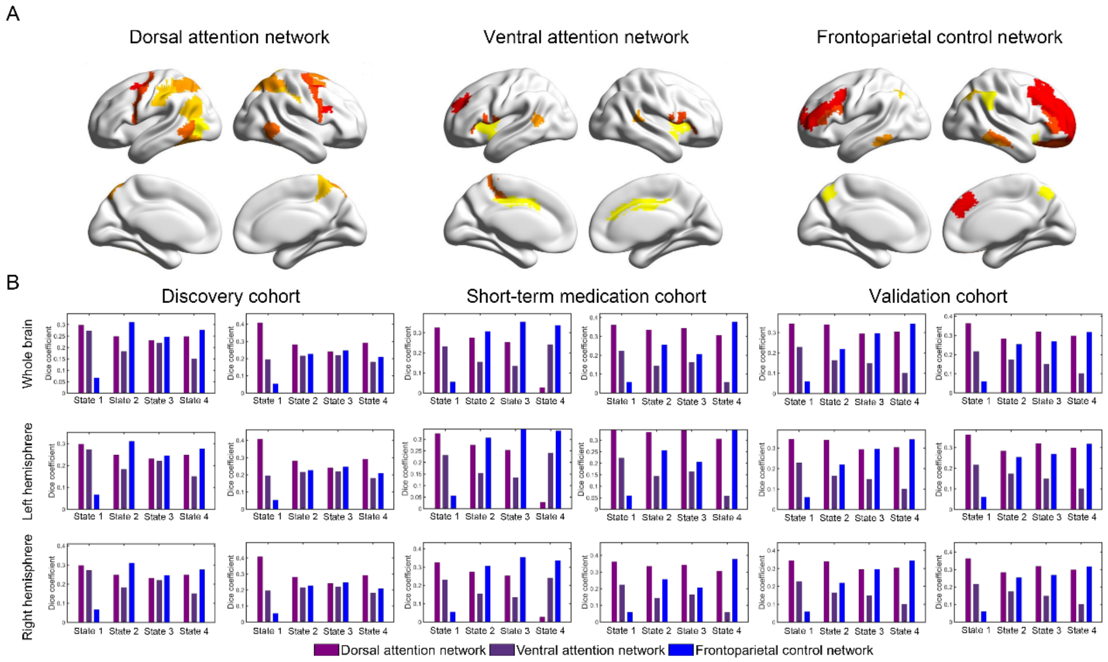


**Supplementary Figure 3. Functional relevance of hubs in the first three states of the executive control network.** A. The parcels corresponding to the dorsal attention network, ventral attention network, and frontoparietal network were defined from Brainnetome Atlas (https://atlas.brainnetome.org/bnatlas.html). B: Dice coefficients between binary images of hubs and meta-analysis results. The hubs in State 1 primarily originated from the dorsal and ventral attention networks, those in State 2 mainly from the dorsal attention and frontoparietal control networks, while the hubs in State 3 showed a comparable degree of overlap among the three networks.

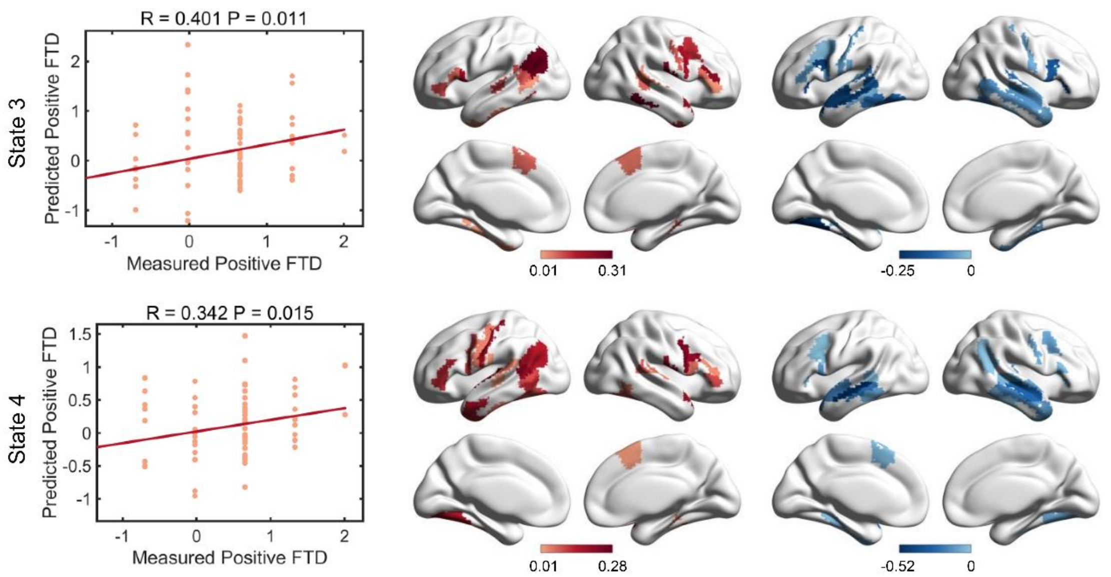


**Supplementary Figure 4. Model prediction and significance of positive FTD by the language network in the discovery cohort**. In each model, nodes with top 5% (red) of positive weights and nodes with top 5% (blue) of negative weights were visualized.

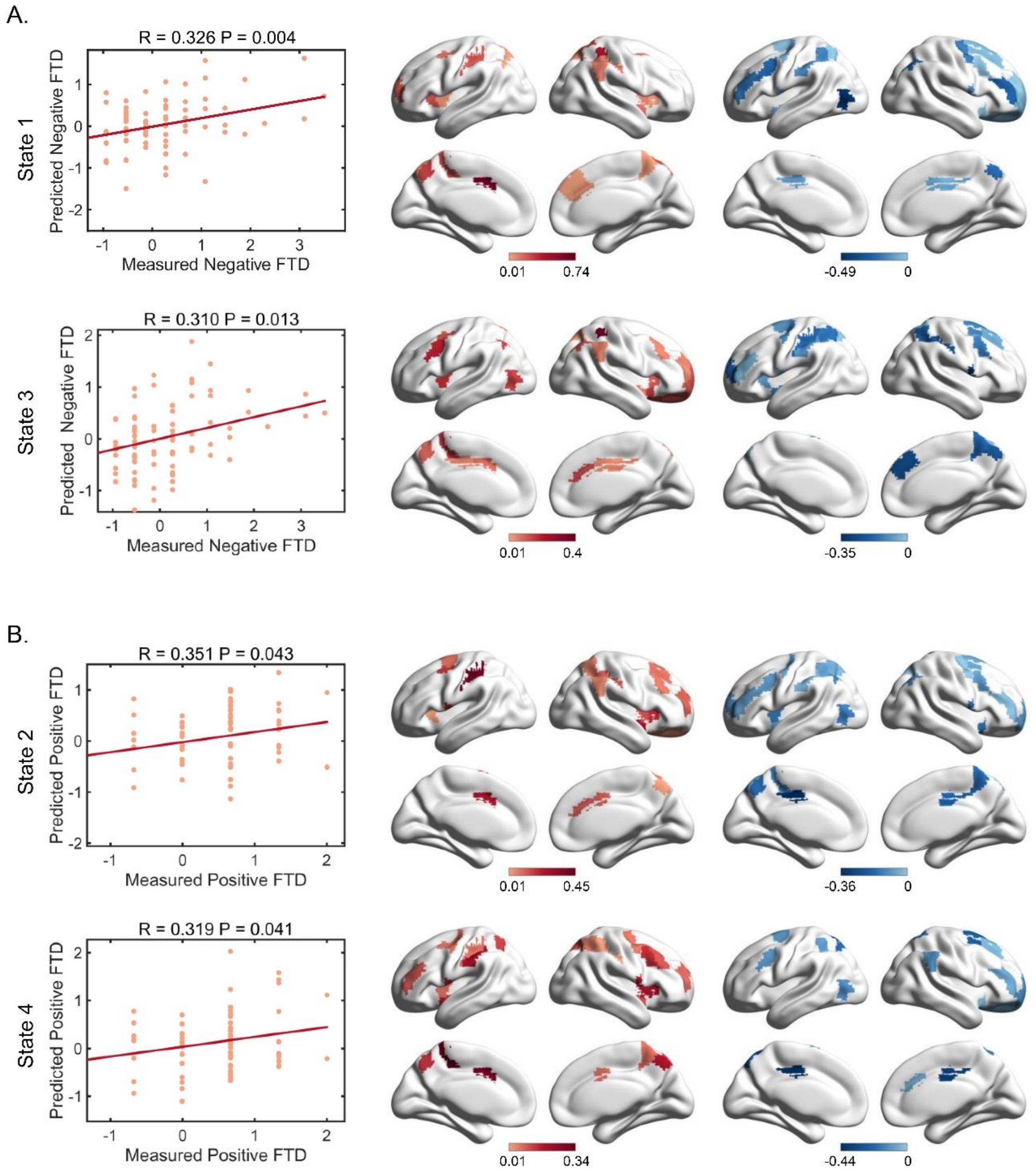


**Supplementary Figure 5. Model prediction and significance of FTD by the executive control network in the discovery cohort**. A. Model prediction and significance of negative FTD by the executive control network. In each model, nodes with top 5% (red) of positive weights and nodes with top 5% (blue) of negative weights were visualized. B. Model prediction and significance of positive FTD by the executive control network.

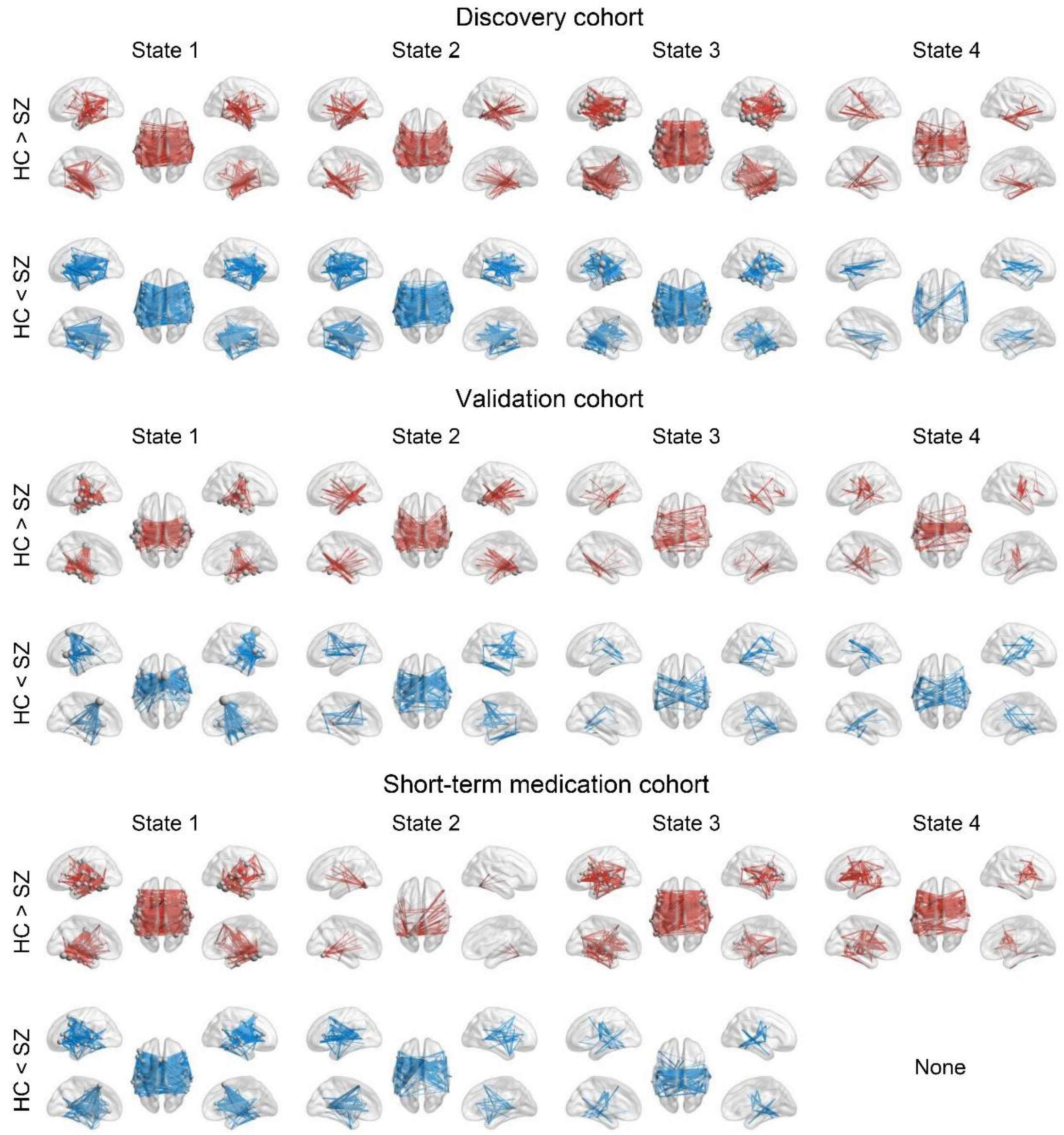


**Supplementary Figure 6. The state-specific hyper- and hypo-connectivity of language network in individuals with schizophrenia.** For each state, between-group comparison was corrected using NBS with an edge *P* value of 0.01 and a component *P* value of 0.01. HC, healthy control; SZ, schizophrenia.

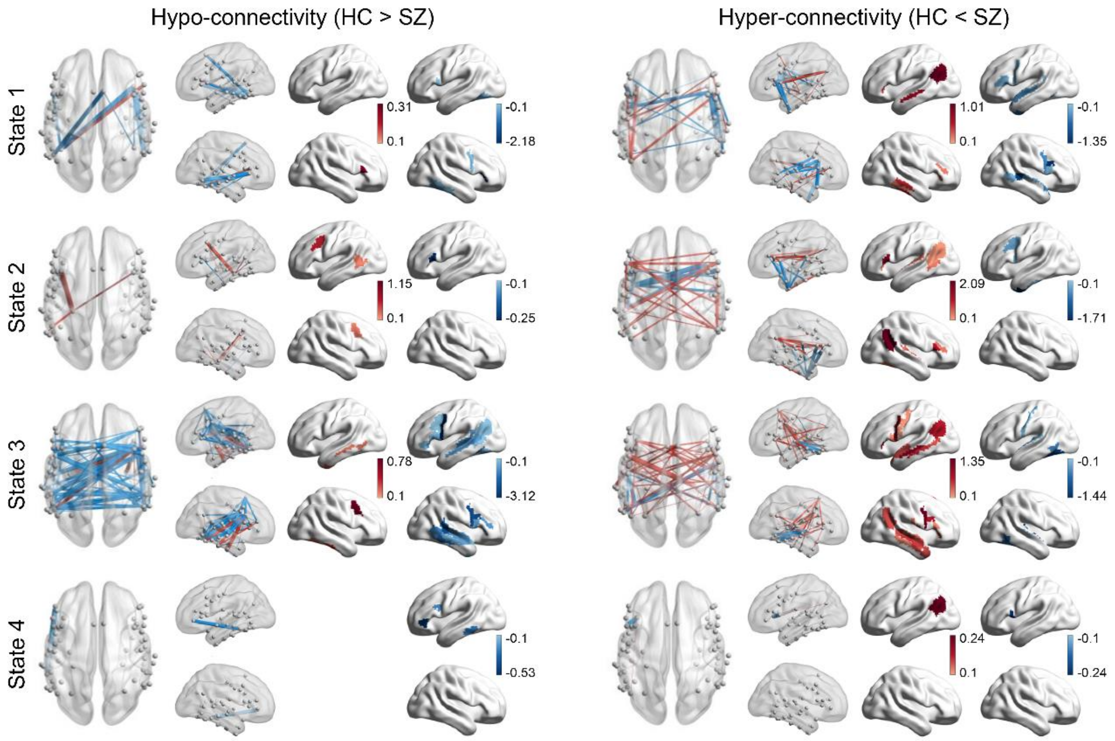


**Supplementary Figure 7. Hypo- and hyper-connectivity of the language network and their associations with positive FTD in the discovery cohort.** The blue lines indicate a negative correlation with positive FTD, while the red lines indicate a positive correlation. Pearson correlation coefficients were calculated using sex, age, and head motion as covariates. Only edges with a significant correlation ($P < 0.05$ with NBS correction) with behavioral data were retained.

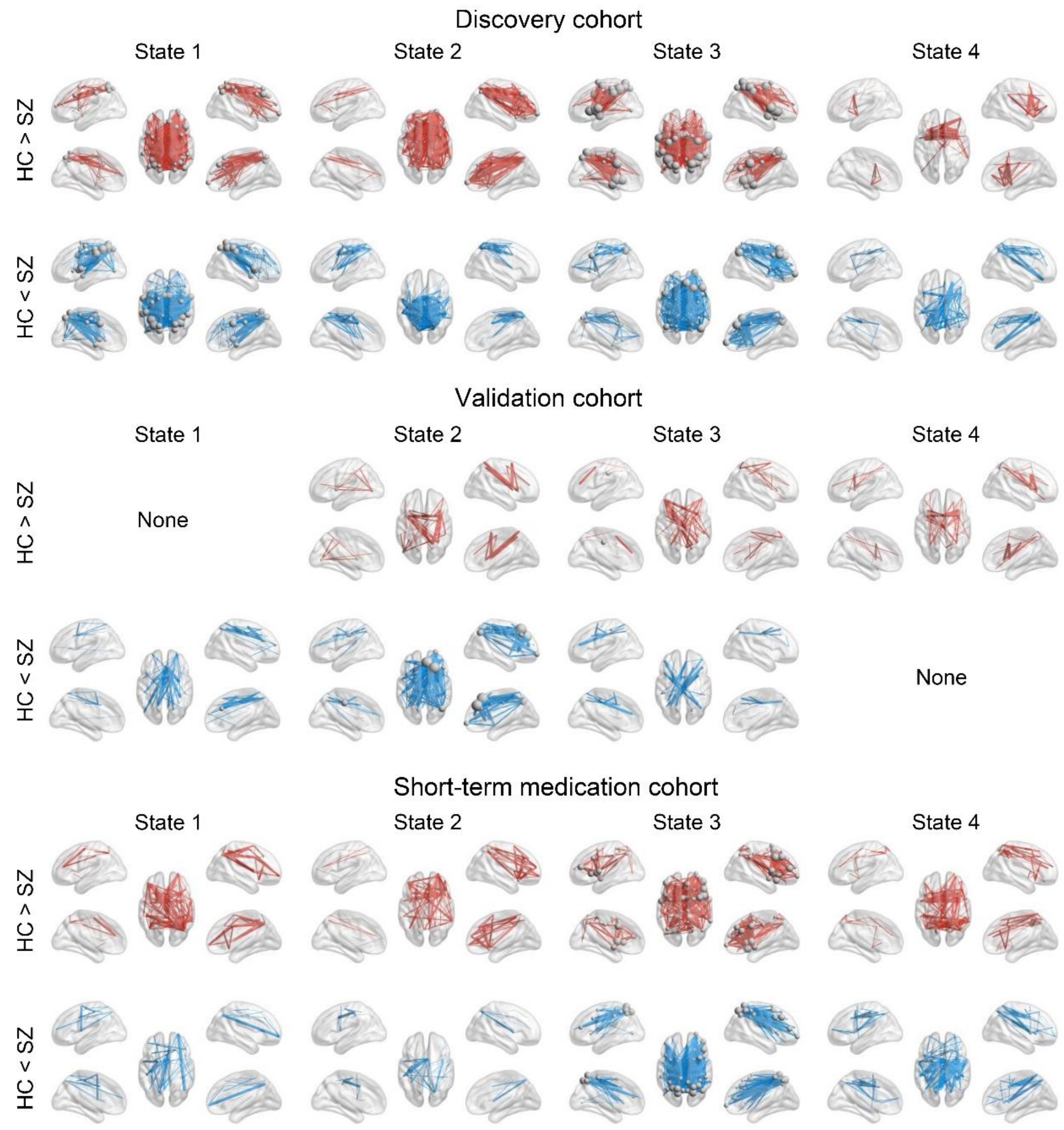


**Supplementary Figure 8. The state-specific hyper- and hypo-connectivity of the executive control network in individuals with schizophrenia.** For each state, between-group differences were corrected using NBS with an edge *P* value of 0.01 and a component *P* value of 0.01.

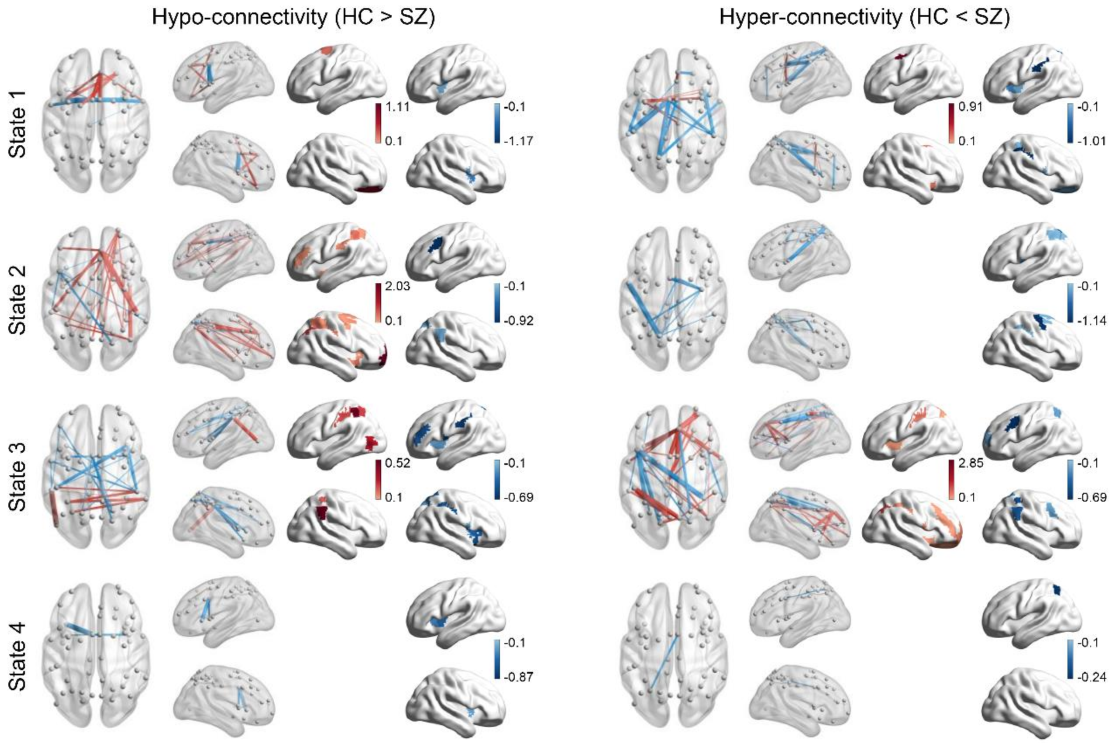


**Supplementary Figure 9. Hypo- and hyper-connectivity of the executive control network and their associations with positive FTD in the discovery cohort.**

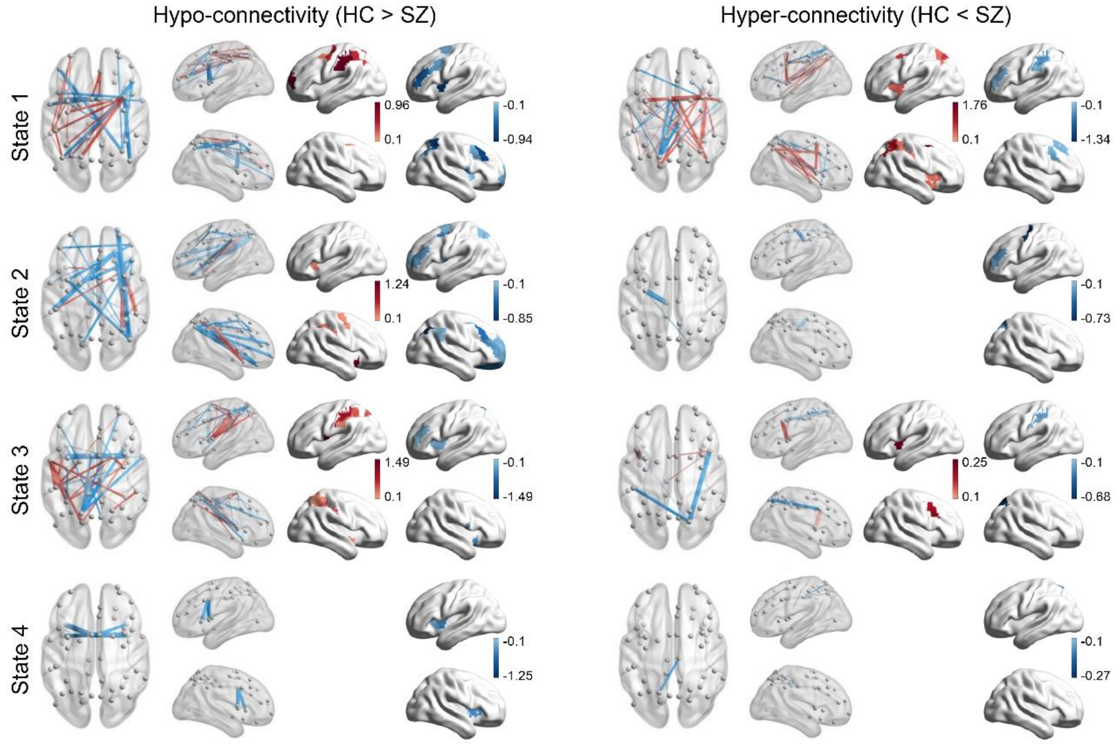


**Supplementary Figure 10. Hypo- and hyper-connectivity of the executive control network and their associations with negative FTD in the discovery cohort.**

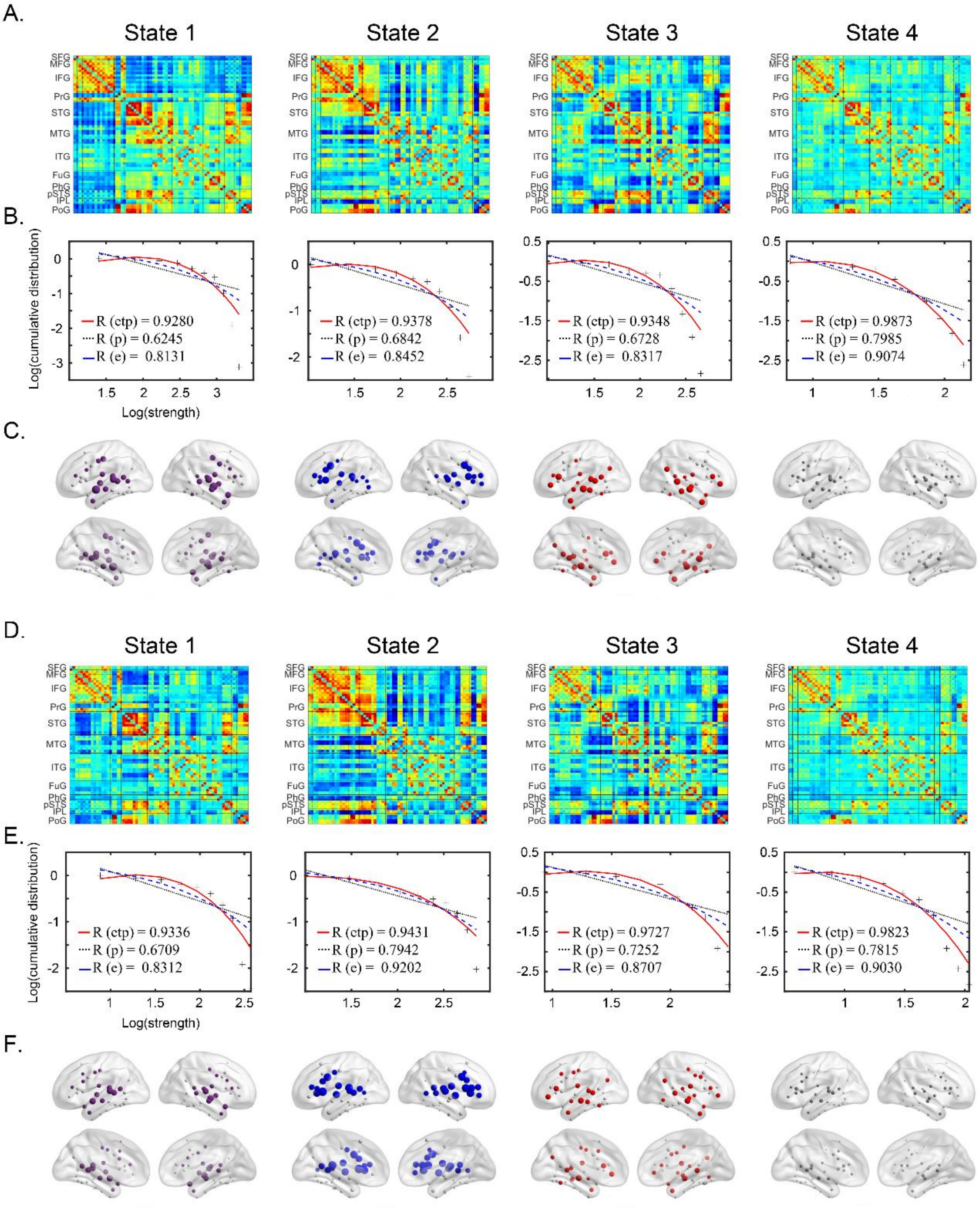


**Supplementary Figure 11. The four meta-networking states of the language network in the validation cohort.** (A & C). Connectivity matrices of the four states in HCs and individuals with schizophrenia. In each group, four temporally reoccurring states with distinct functional connectivity patterns were observed. (B & D). Log-log plots of the cumulative nodal strength distributions. The plus sign (black) represents observed data, the solid line (red) represents the fit of the exponentially truncated power-law distribution, $P(x) \sim x^{\alpha-1}\exp(\frac{x}{x_c})$, the dashed line (blue) represents an exponential distribution, $P(x) \sim \exp(\frac{x}{x_c})$, and the dotted line (black) represents a power-law, $P(x) \sim x^{\alpha-1}$. $R^2$ was calculated to assess the goodness of fit, with larger values indicating better fitting. The exponentially truncated power-law is the best fitting for all four states. (C & F). Nodal strength distributions. The first 20 nodes with the highest nodal strength were defined as hubs. State-dependent hub distributions were observed.

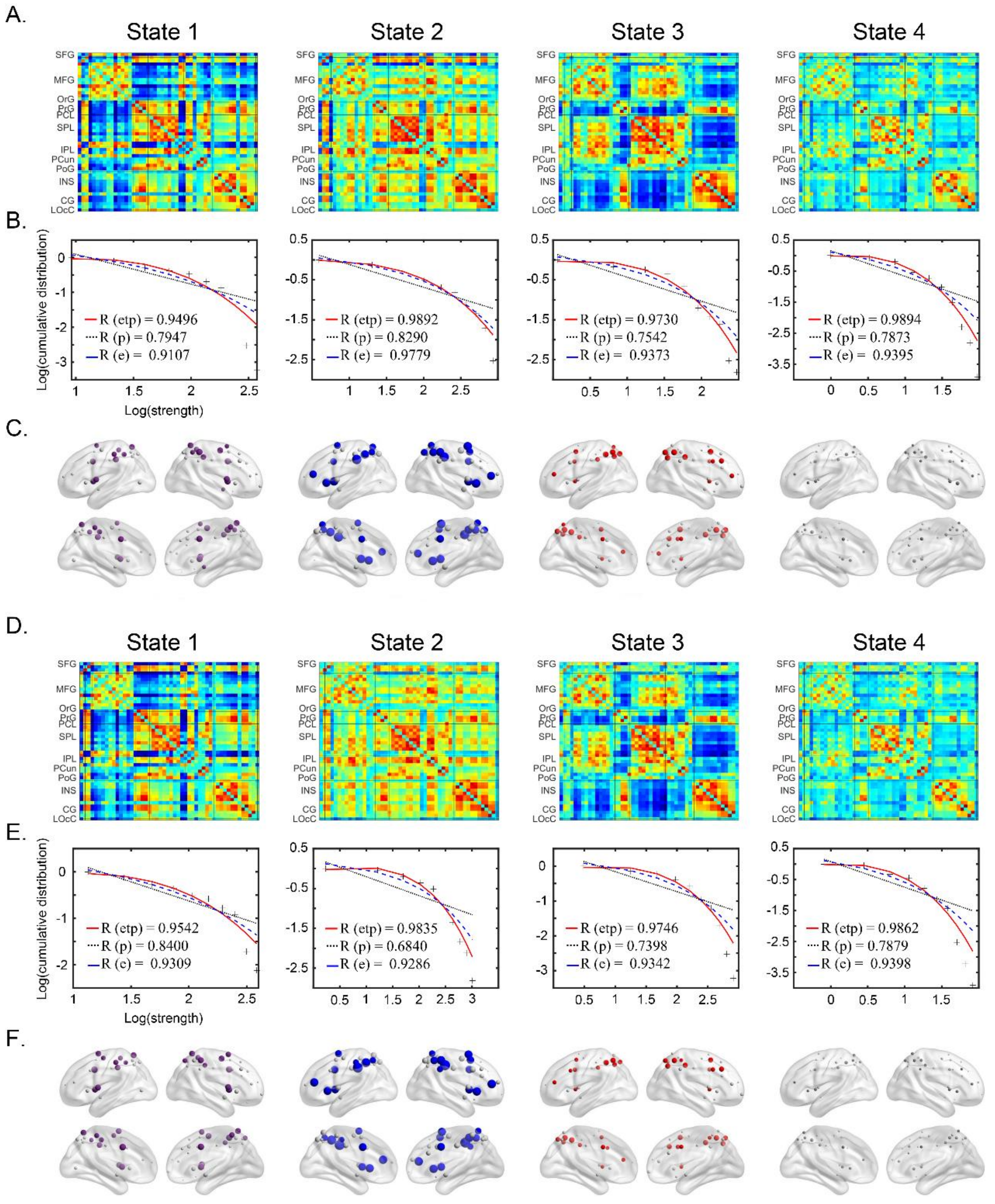


**Supplementary Figure 12. The four meta-networking states of the executive control network in the validation cohort.**

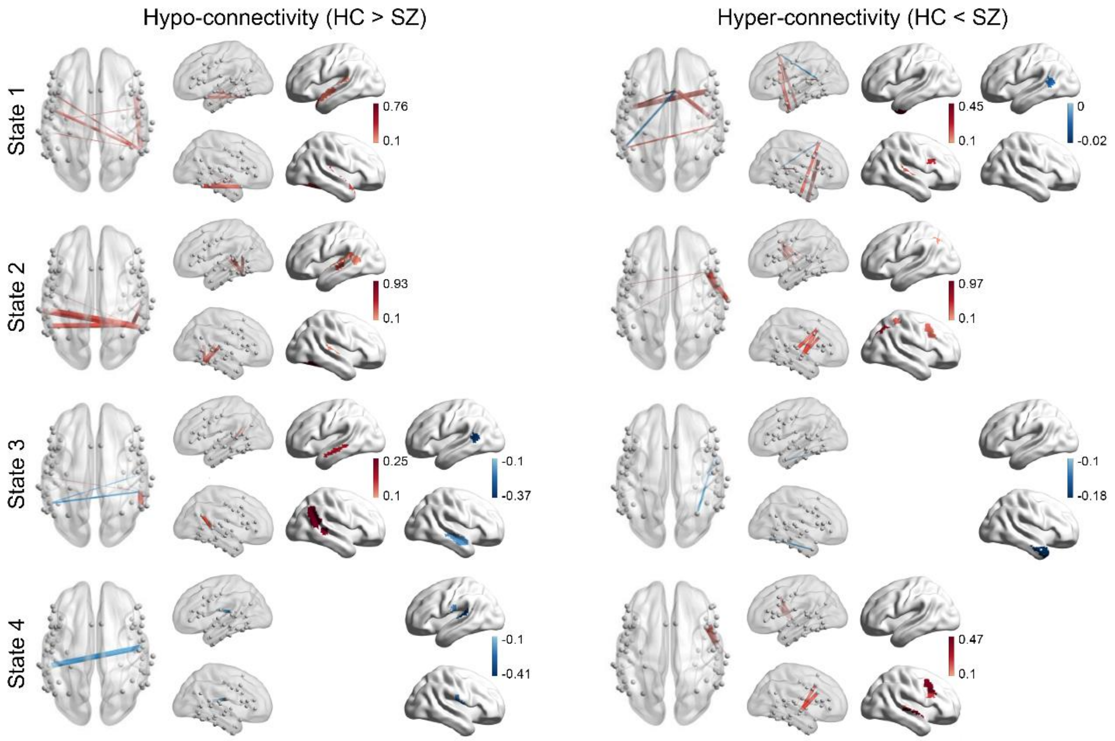


**Supplementary Figure 13. Hypo- and hyper-connectivity of the language network and their associations with positive FTD in the validation cohort.**

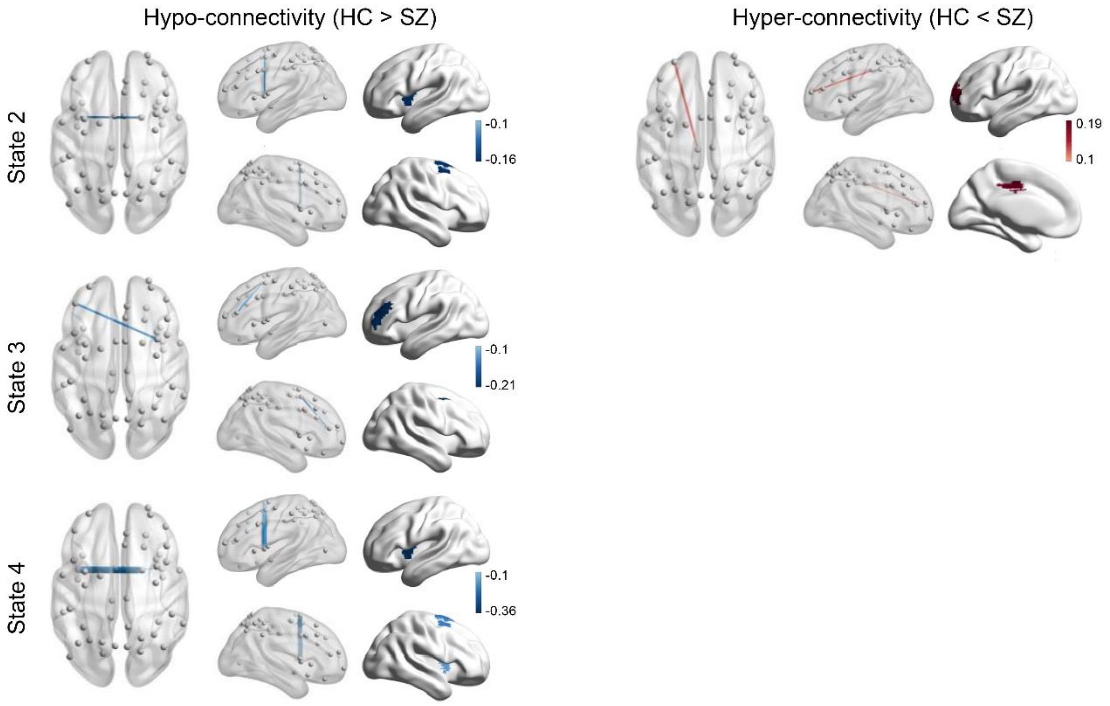


**Supplementary Figure 14. Hypo- and hyper-connectivity of the executive control network and their associations with negative FTD in the validation cohort.**

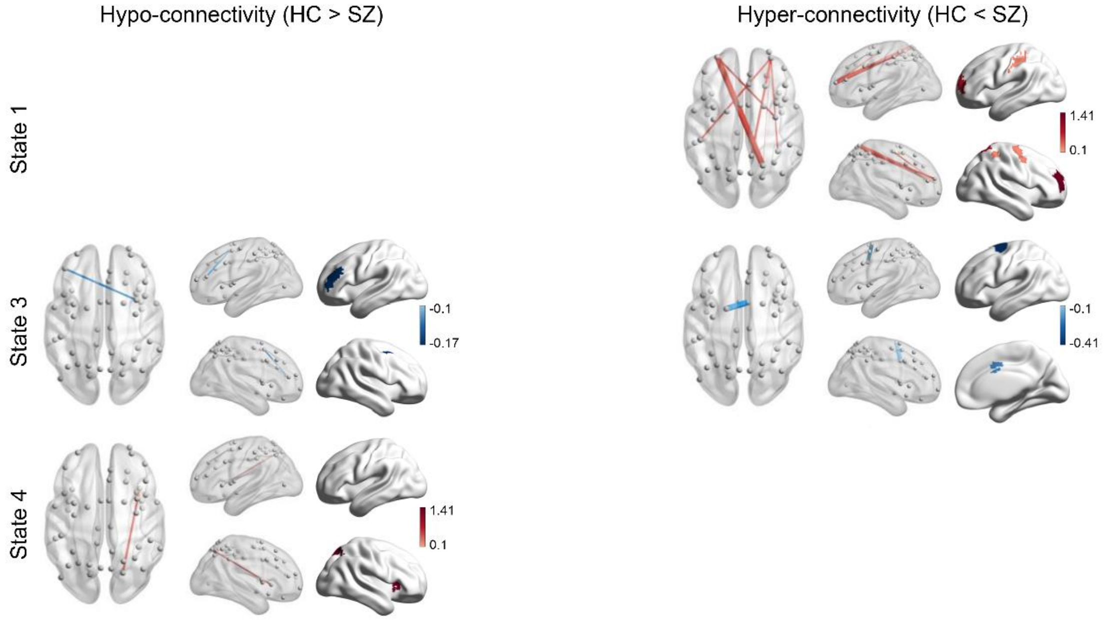


**Supplementary Figure 15. Hypo- and hyper-connectivity of the executive control network and their associations with positive FTD in the validation cohort.**

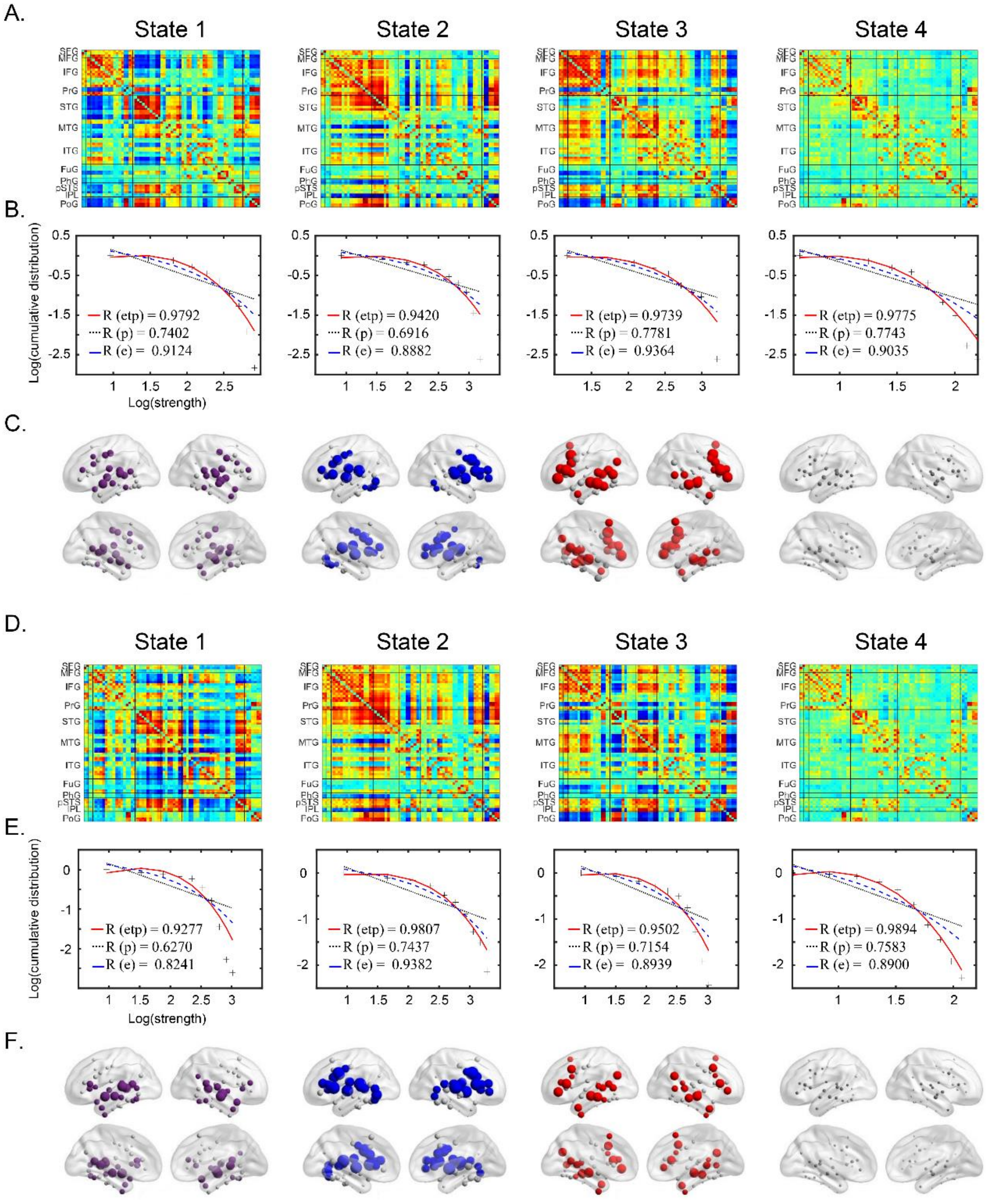


**Supplementary Figure 16. The four meta-networking states of the language network in the short-term medication cohort.**

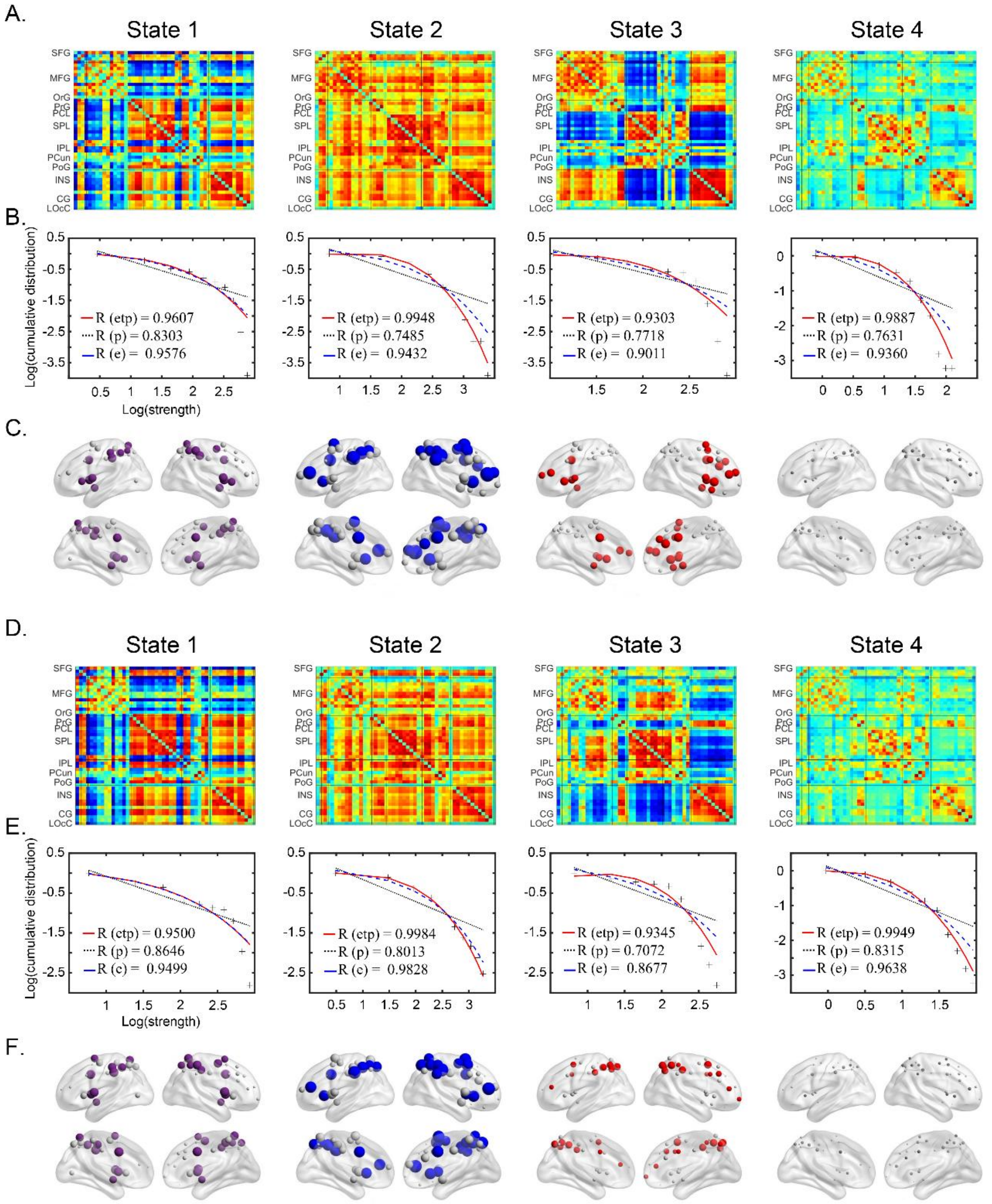


**Supplementary Figure 17. The four meta-networking states of the executive control network in the short-term medication cohort.**

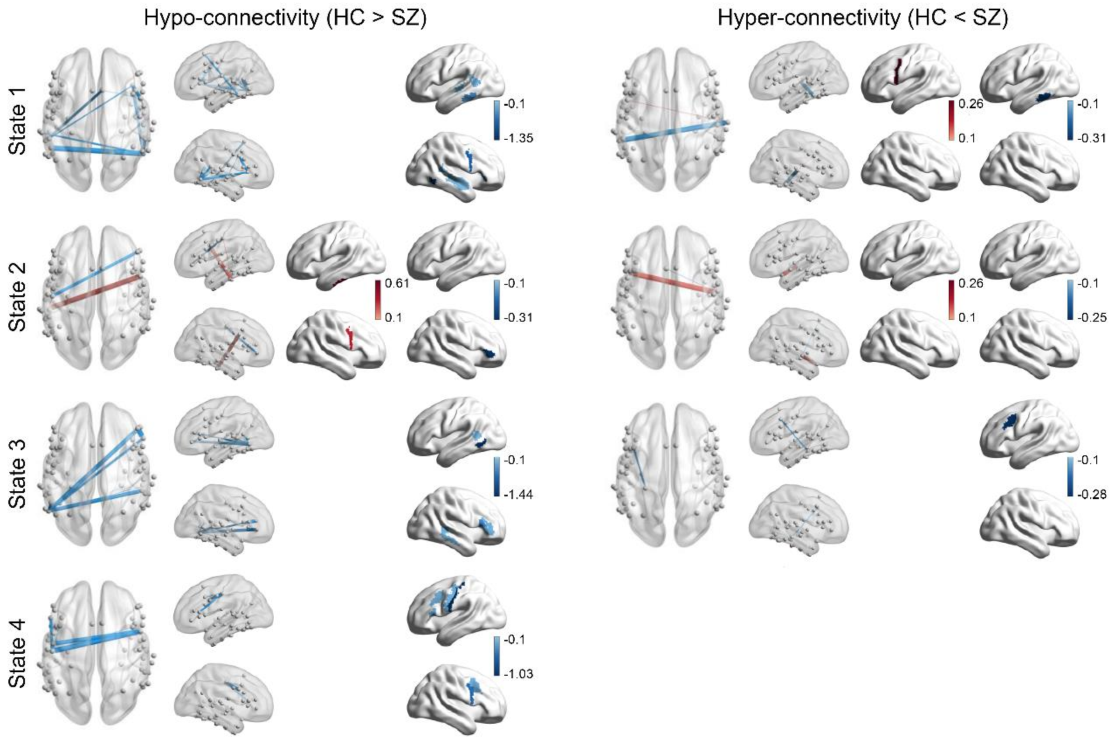


**Supplementary Figure 18. Hypo- and hyper-connectivity of the language network and their associations with positive FTD in the short-term medication cohort.**

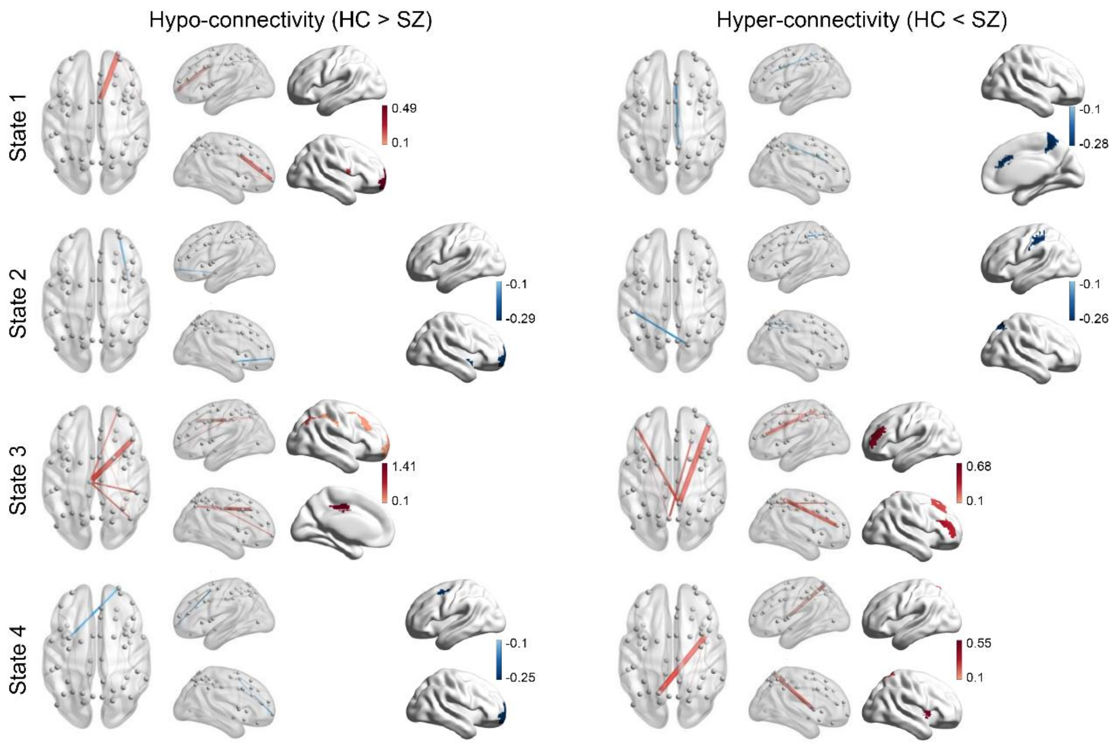


**Supplementary Figure 19. Hypo- and hyper-connectivity of the executive control network and their associations with positive FTD in the short-term medication cohort.**

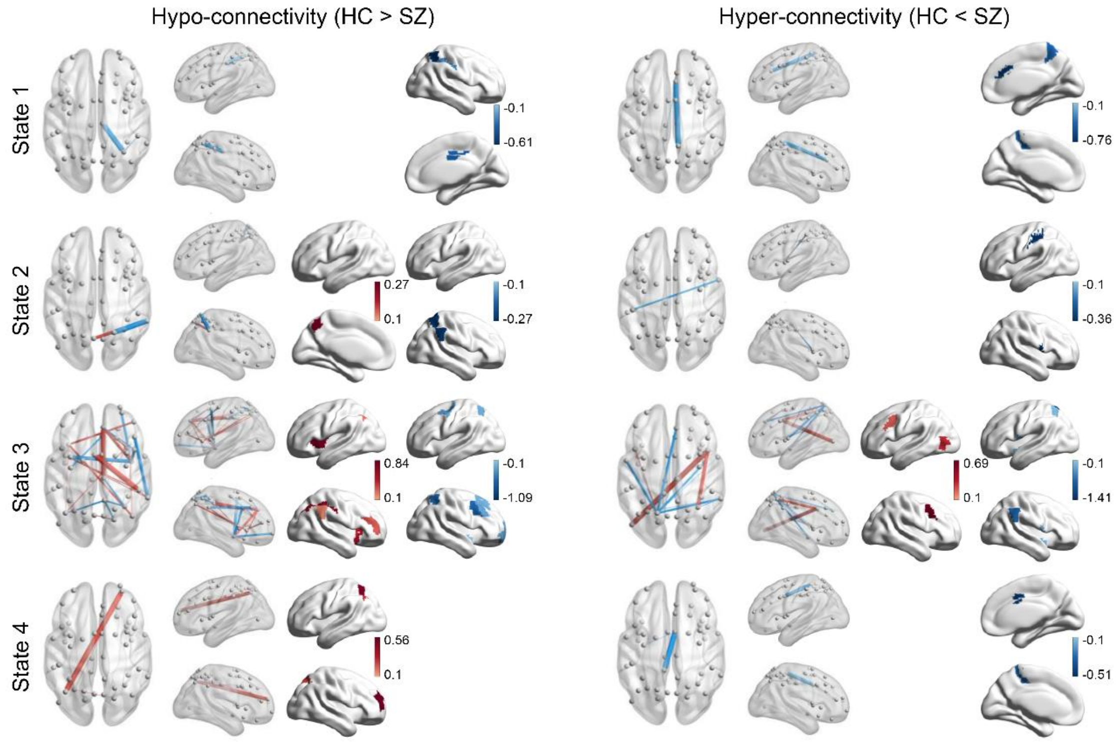


**Supplementary Figure 20. Hypo- and hyper-connectivity of the executive control network and their associations with negative FTD in the short-term medication cohort.**

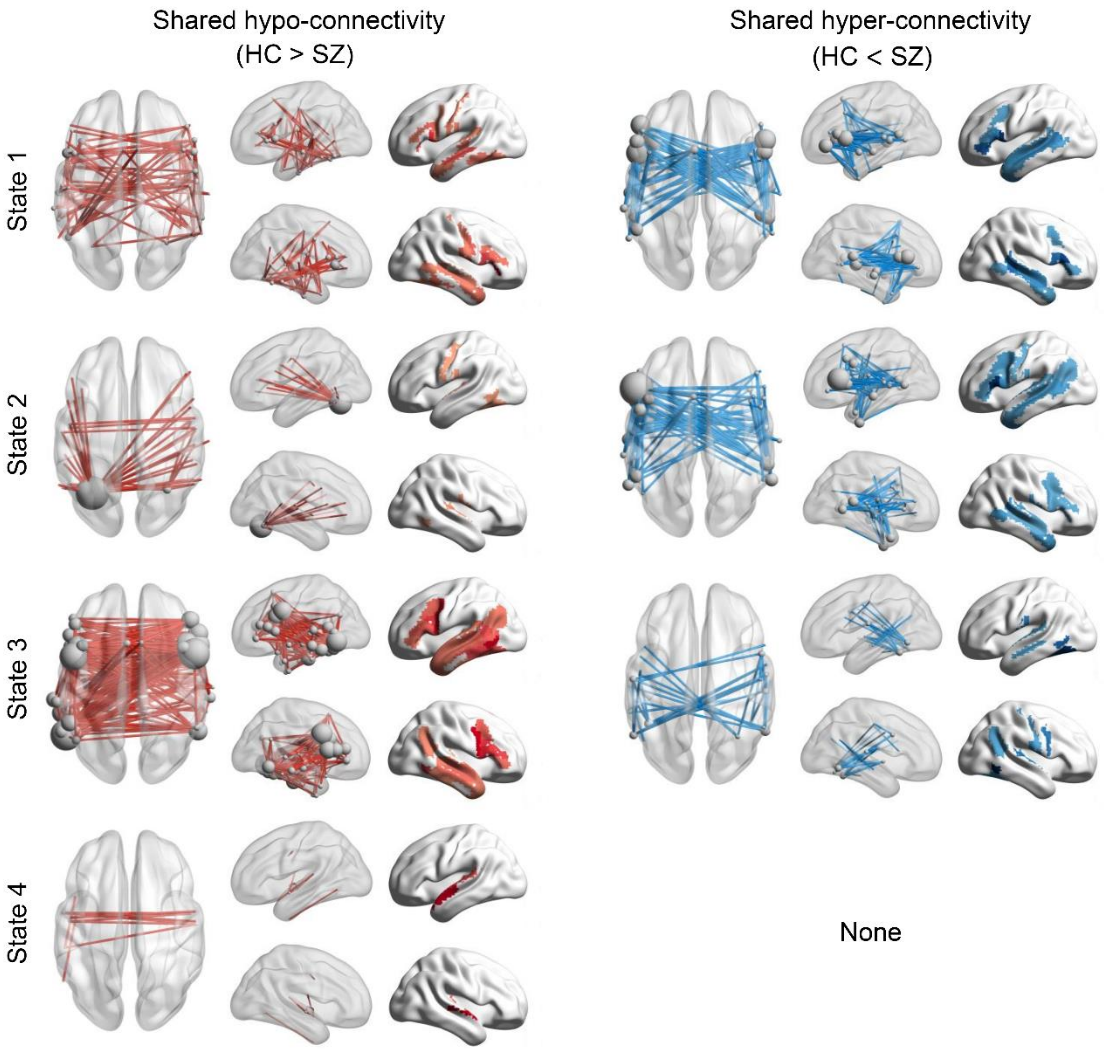


**Supplementary Figure 21. Between-site shared hypo- and hyper-connectivity of language network in schizophrenia**. Shared hypo- and hyper-connectivity in each state of the language network.

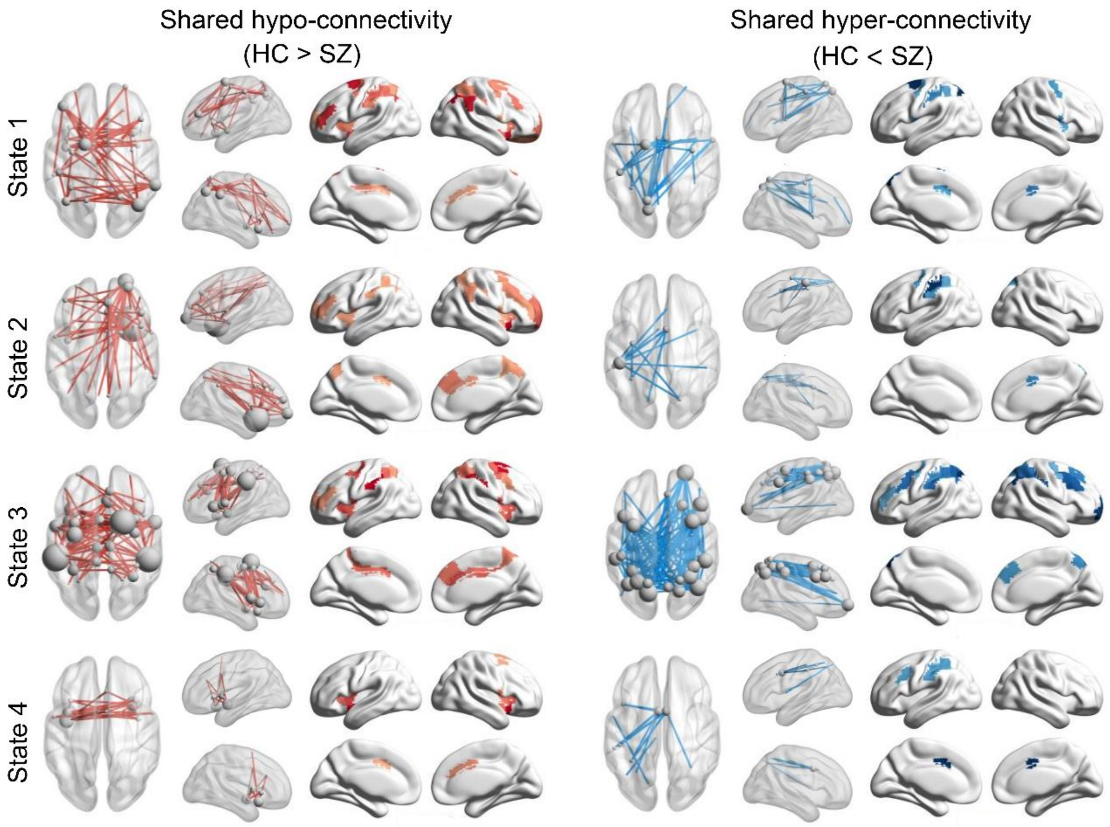


**Supplementary Figure 22. Between-site shared hypo- and hyper-connectivity of executive control network in schizophrenia**. Shared hypo- and hyper-connectivity in each state of the executive control network.

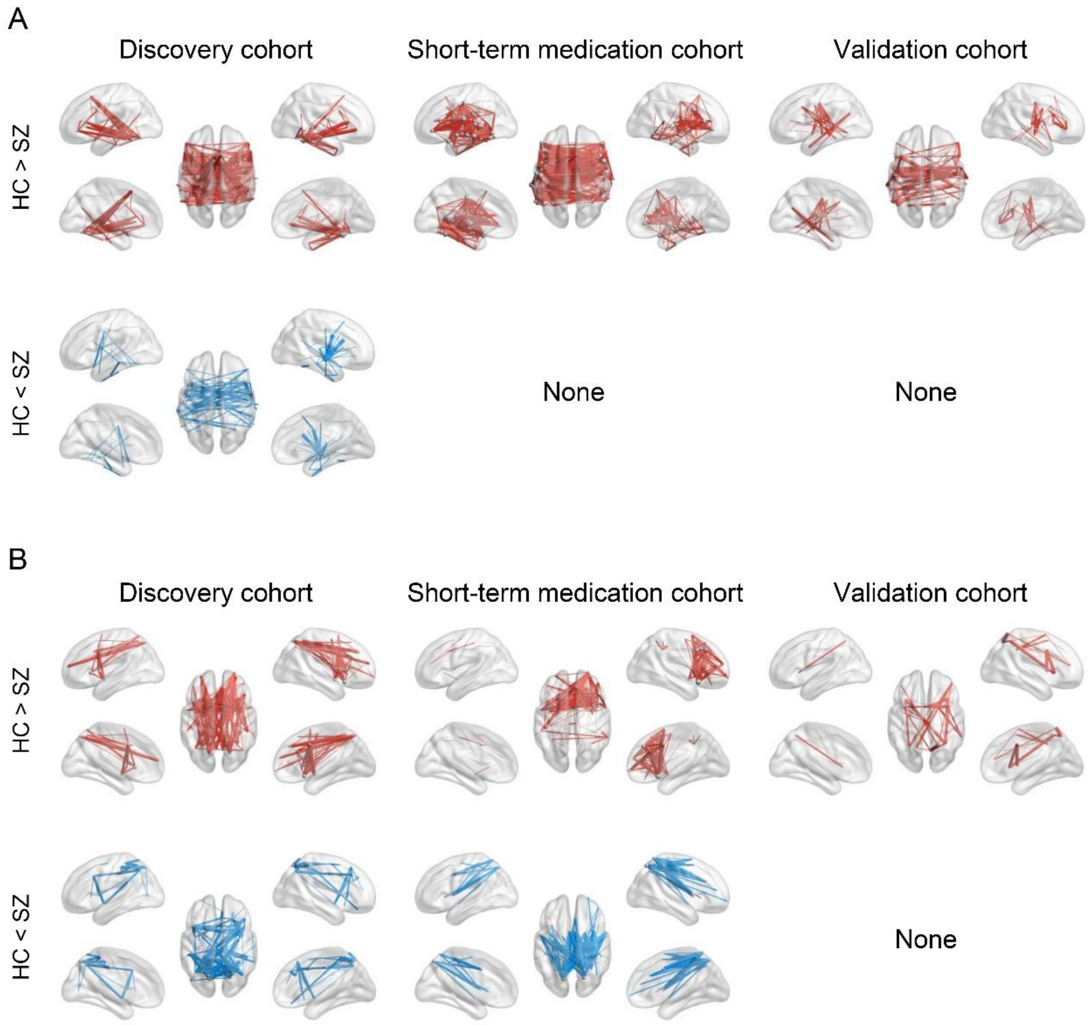


**Supplementary Figure 23. Group differences of sFC of the executive control network.** Edge $P < 0.01$, Component $P < 0.01$, NBS correction.

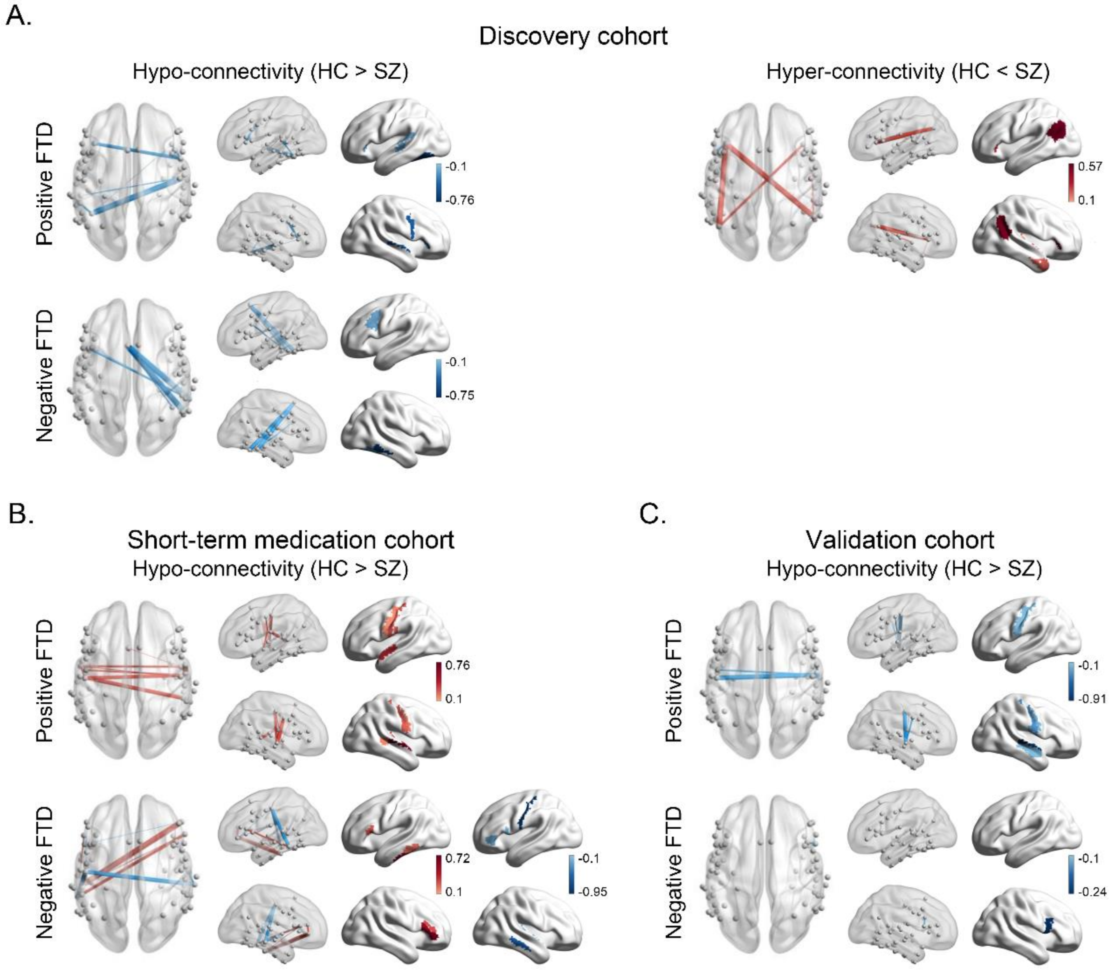


**Supplementary Figure 24. Partial correlations between FTD scores and sFC in the language network**.

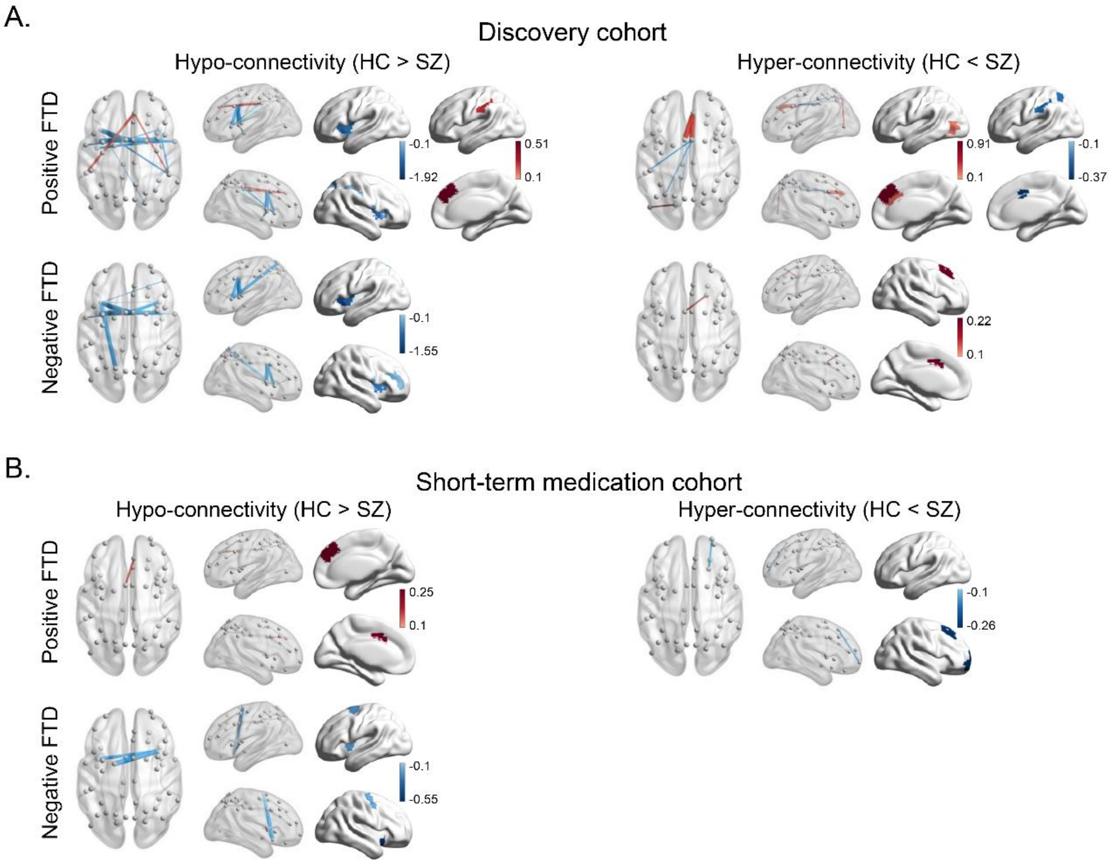


**Supplementary Figure 25. Partial correlations between FTD scores and sFC in the executive control network**.

#### Supplementary tables

**Supplementary Table 1. Anatomical regions，language-related behavioral domains, and paradigm classes of the language network. Coordinates are in the standard Montreal Neurologic Institute Space.**

| ID | x | y | z | anatomy | behavioral domains | paradigm classes |
|---|---|---|---|---|---|---|
| 1 | -4.6 | 15.7 | 53.2 | SFG | Execution. Speech, Phonology, Semantics, Speech | Word Generation (Covert and Overt) |
| 2 | 7.0 | 16.4 | 54.7 | SFG | The homolog of node 1 | - |
| 17 | -41.9 | 13.6 | 36.4 | MFG | Phonology, Semantics | Semantic. Monitor/Discrimination, Word Generation (Covert) |
| 18 | 42.2 | 11.9 | 38.4 | MFG | The homolog of node 17 | - |
| 29 | -45.6 | 13.0 | 23.4 | IFG | Phonology, Semantics, Speech, and Syntax | Phonological. Discrimination, Semantic. Monitor/Discrimination |
| 30 | 45.0 | 15.7 | 25.0 | IFG | The homolog of node 29 | - |
| 31 | -47.6 | 31.6 | 13.6 | IFG | Phonology, Semantics, Speech, and Syntax | Phonological. Discrimination, Semantic. Monitor/Discrimination, Word Generation (Covert and Overt) |
| 32 | 47.8 | 35.1 | 13.2 | IFG | The homolog of node 31 | - |
| 33 | -52.2 | 22.5 | 11.4 | IFG | Semantics, Speech, and Syntax | Reading (Covert), Semantic. Monitor/Discrimination, Word Generation (Covert and Overt) |
| 34 | 54.2 | 23.9 | 11.8 | IFG | The homolog of node 33 | - |
| 35 | -49.1 | 36.3 | -2.8 | IFG | Semantics, Speech, and Syntax | Semantic. Monitor/Discrimination, Word Generation (Covert) |
| 36 | 50.8 | 36.8 | -0.7 | IFG | The homolog of node 35 | - |
| 37 | -39.5 | 22.9 | 3.7 | IFG | Phonology, Semantics, Speech, and Syntax | Semantic. Monitor/Discrimination, Word Generation (Covert) |
| 38 | 42.1 | 22.0 | 3.2 | IFG | The homolog of node 37 | - |
| 39 | -51.3 | 13.2 | 6.0 | IFG | Phonology, Semantics, Speech | Music. Comprehension/Production, Recitation/Repetition. (Covert), Word Generation (Covert) |
| 40 | 53.6 | 14.3 | 11.8 | IFG | The homolog of node 39 | - |
| 53 | -49.5 | -7.1 | 38.8 | PrG | Execution. Speech | Reading (Overt), Recitation/Repetition. (Overt) |
| 54 | 54.9 | -2.0 | 33.3 | PrG | Execution. Speech | Reading (Overt), Recitation/Repetition. |

| | | | | | | |
|---|---|---|---|---|---|---|
| | | | | | | (Overt) |
| 63 | -49.1 | 4.7 | 30.5 | PrG | Orthography, Phonology, Semantics, Speech, and Syntax | Phonological. Discrimination, Reading (Covert) |
| 64 | 51.1 | 7.2 | 30.9 | PrG | The homolog of node 63 | - |
| 71 | -53.7 | -32.1 | 12.4 | STG | Execution. Speech, Phonology, and Speech | Music. Comprehension/Production, Passive Listening, Phonological. Discrimination, Reading (Overt), Recitation/Repetition. (Covert and Overt) |
| 72 | 54.5 | -23.7 | 10.6 | STG | Execution. Speech, Phonology | Music. Comprehension/Production, Passive Listening, Phonological. Discrimination, Recitation/Repetition. (Overt) |
| 73 | -50.1 | -10.3 | 1.1 | STG | Execution. Speech, Phonology, Speech | Music. Comprehension/Production, Passive Listening, Phonological. Discrimination, Reading (Overt), Recitation/Repetition. (Overt) |
| 74 | 51.1 | -3.7 | -0.9 | STG | Execution. Speech, | Music. Comprehension/Production, Passive Listening, Recitation/Repetition. (Overt) |
| 75 | -62.8 | -32.8 | 7.4 | STG | Execution. Speech, Phonology, Semantics, Speech | Passive Listening, Phonological. Discrimination, Reading (Overt), Semantic. Monitor/Discrimination |
| 76 | 66.5 | -20.8 | 6.6 | STG | Execution. Speech, Phonology, Speech | Music. Comprehension/Production, Passive Listening, Phonological. Discrimination, Reading (Overt) |
| 77 | -45.1 | 10.5 | -19.4 | STG | The homolog of node 78 | - |
| 78 | 47.1 | 12.3 | -19.7 | STG | Speech | Film Viewing, Passive Listening |
| 79 | -55.1 | -3.2 | -10.1 | STG | Execution. Speech, Phonology, Semantics, Speech | Music. Comprehension/Production, Passive Listening, Phonological. Discrimination, Reading (Overt) |
| 80 | 55.8 | -12.5 | -5.2 | STG | Execution. Speech, Phonology, Semantics, Speech | Music. Comprehension/Production, Passive Listening, Phonological. Discrimination, Reading (Overt), Semantic. Monitor/Discrimination |
| 81 | -65.2 | -30.9 | -11.3 | MTG | Semantics | Semantic. Monitor/Discrimination |
| 82 | 64.5 | -29.2 | -13.2 | MTG | The homolog of node 81 | - |
| 83 | -53.2 | 2.2 | -29.6 | MTG | Language | Semantic. Monitor/Discrimination, Reading (Covert) |
| 84 | 51.1 | 5.7 | -31.8 | MTG | Language | Passive Listening |
| 85 | -58.9 | -57.6 | 4.3 | MTG | Semantics and Syntax | Semantic. Monitor/Discrimination, Word Generation (Overt) |
| 86 | 60.1 | -53.3 | 2.9 | MTG | The homolog of node | Film Viewing |

| | | | | | | |
|---|---|---|---|---|---|---|
| | | | | | 85 | |
| 87 | -58.5 | -19.8 | -9.4 | MTG | Phonology, Semantics, Speech, and Syntax | Passive Listening, Phonological. Discrimination, Reading (Covert), Semantic. Monitor/Discrimination |
| 88 | 58.3 | -15.4 | -10.1 | MTG | Semantics and Speech | Passive Listening, Phonological. Discrimination |
| 89 | -45.5 | -26.7 | -26.1 | ITG | Orthography and Semantics | Reading (Covert), Semantic. Monitor/Discrimination |
| 90 | 45.8 | -14.6 | -32.4 | ITG | The homolog of node 89 | - |
| 91 | -50.5 | -57.0 | -14.1 | ITG | Phonology and Semantics | Naming (Overt) |
| 92 | 53.5 | -52.4 | -18.5 | ITG | Phonology and Semantics | - |
| 93 | -43.7 | -2.9 | -41.4 | ITG | Semantics | Semantic. Monitor/Discrimination |
| 94 | 40.5 | -2.9 | -41.4 | ITG | The homolog of node 93 | - |
| 97 | -55.2 | -60.3 | -6.0 | ITG | Semantics and Speech | Film Viewing, Naming (Overt) |
| 98 | 54.2 | -56.9 | -8.6 | ITG | The homolog of node 97 | - |
| 99 | -58.8 | -42.1 | -16.0 | ITG | Orthography and Semantics | Naming (Overt), Reading (Covert), Semantic. Monitor/Discrimination, Word Generation (Overt) |
| 100 | 60.5 | -40.5 | -17.1 | ITG | The homolog of node 99 | - |
| 101 | -54.9 | -30.3 | -27.4 | ITG | Semantics | - |
| 102 | 53.7 | -30.3 | -26.3 | ITG | The homolog of node 101 | - |
| 103 | -32.4 | -16.6 | -32.3 | FuG | Semantics and Speech | Naming (Overt), Semantic. Monitor/Discrimination |
| 104 | 33.1 | -14.6 | -34.1 | FuG | Semantics | Naming (Overt), Semantic. Monitor/Discrimination |
| 105 | -30.6 | -64.4 | -14.1 | FuG | Orthography, Semantics, Speech | Naming (Covert and Overt) |
| 106 | 31.3 | -61.4 | -13.7 | FuG | Language | Naming (Covert and Overt) |
| 107 | -42.3 | -50.9 | -17.3 | FuG | Orthography, Phonology, Semantics, Speech | Naming (Covert and Overt), Phonological. Discrimination, Reading (Covert), and Semantic. Monitor/Discrimination |
| 108 | 42.7 | -49.1 | -18.6 | FuG | Orthography, Semantics | Naming (Covert) |
| 113 | -28.3 | -32.6 | -16.9 | PhG | Semantics | Naming (Overt), Semantic. Monitor/Discrimination |
| 114 | 28.8 | -30.7 | -17.5 | PhG | The homolog of node | Passive Listening, Semantic. |

| | | | | | | |
|---|---|---|---|---|---|---|
| | | | | | 113 | Monitor/Discrimination |
| 121 | -54.4 | -39.8 | 4.2 | pSTS | Phonology, Semantics, Speech, Syntax | Passive Listening, Phonological. Discrimination, Reading (Covert), Semantic. Monitor/Discrimination, Word Generation (Covert and Overt) |
| 122 | 52.9 | -36.8 | 3.1 | pSTS | Execution. Speech, Phonology, Semantics, and Speech | Passive Listening, Phonological. Discrimination, Reading (Overt) |
| 123 | -52.4 | -50.3 | 10.8 | pSTS | Orthography, Semantics, Speech, and Syntax | Reading (Covert), Semantic. Monitor/Discrimination |
| 124 | 56.5 | -40.1 | 12.5 | pSTS | The homolog of node 123 | Passive Listening |
| 143 | -46.8 | -64.7 | 25.8 | IPL | Language | Semantic. Monitor/Discrimination, |
| 144 | 53.0 | -54.1 | 24.4 | IPL | The homolog of node 143 | - |
| 155 | -50.4 | -15.8 | 42.1 | PoG | Execution. Speech | Recitation/Repetition. (Overt) |
| 156 | 50.3 | -14.2 | 43.7 | PoG | Execution. Speech | Reading (Overt), Recitation/Repetition. (Overt) |
| 157 | -55.8 | -14.0 | 16.2 | PoG | Execution. Speech | Recitation/Repetition. (Overt) |
| 158 | 55.2 | -10.2 | 15.0 | PoG | Execution. Speech | Recitation/Repetition. (Overt) |

*SFG*, superior frontal gyrus; *MFG*, middle frontal gyrus; *IFG*, inferior frontal gyrus; *OrG*, orbital gyrus; *PrG*, precentral gyrus; *STG*, superior temporal gyrus; *MTG*, middle temporal gyrus; *ITG*, inferior temporal gyrus; *FuG*, fusiform gyrus; *PhG*, hippocampal gyrus; *pSTS*, posterior superior temporal sulcus; *IPL*, inferior parietal lobule. *PoG*, postcentral gyrus. The ID is the number of the parcel in the Brainnetome atlas (Fan et al., 2016). Here we only summarized the language-related behavioral domains and paradigm classes; the full behavioral domains and paradigm classes for each node are available at http://atlas.brainnetome.org/bnatlas.php.

**Supplementary Table 2. Anatomical regions, executive control-related behavioral domains, and paradigm classes of the executive control network. Coordinates are in the standard Montreal Neurologic Institute Space.**

| ID | x | y | z | anatomy | behavioral domains | paradigm classes |
|---|---|---|---|---|---|---|
| 4 | 21.9 | 25.7 | 51.1 | SFG | Social Cognition | Theory of Mind Task |
| 7 | -17.9 | -0.5 | 65.1 | SFG | Vision.Shape, Memory.Working, Motor Learning | Saccades, Mental Rotation, Writing |
| 8 | 19.9 | 4.7 | 63.7 | SFG | Attention, Execution, Memory.Working | Anti-Saccades, Counting/Calculation, Drawing |
| 12 | 5.9 | 38.0 | 35.3 | SFG | Cognition, Inhibition, Emotion, Attention | Reward Task, Stroop Task |
| 17 | -41.9 | 13.6 | 36.4 | MFG | Memory.Explicit, Memory.Working | Encoding, Stroop Task, n-back |
| 18 | 42.2 | 12.0 | 38.3 | MFG | Reasoning, Memory.Working, Inhibition, Attention | Visual Distractor/Visual Attention, n-back, Wisconsin Card Sorting Test |
| 19 | -28.1 | 55.9 | 13.6 | MFG | Inhibition, Social Cognition, Memory.Working | - |
| 20 | 27.8 | 55.0 | 16.6 | MFG | Social Cognition, Memory.Working, Inhibition | Deception Task, Paired Associate Recall, Go/No-Go |
| 21 | -41.0 | 40.6 | 16.4 | MFG | Memory.Working, Reasoning | n-back, Word Generation (Covert), Delayed Match To Sample |
| 22 | 41.5 | 44.3 | 13.6 | MFG | Memory.Working, Perception.Somesthesis .Pain | Delayed Match To Sample, Sternberg Task, Pain.Monitor/Discrimination, n-back |
| 24 | 42.5 | 26.6 | 38.9 | MFG | Memory.Explicit, Reasoning, Inhibition | Flanker Test, n-back, Cued Explicit.Recognition |
| 25 | -31.9 | 3.5 | 54.7 | MFG | Execution, Motor Learning, Vision | n-back, Counting/Calculation, Visual Distractor/Visual Attention |
| 26 | 33.4 | 8.1 | 54.4 | MFG | Preparation, Reasoning, Memory.Working | Mental Rotation, Pointing, n-back, Counting/Calculation, Saccades |
| 28 | 25.6 | 61.2 | -4.0 | MFG | Memory.Explicit | - |
| 46 | 23.4 | 36.5 | -18 | OrG | Gustation, Cognition, Olfaction, Emotion | Reward Task, Olfactory Monitor/Discrimination |
| 55 | -31.8 | -9.0 | 57.6 | PrG | Imagination, Vision.Shape, Execution | Finger Tapping, Mental Rotation, Imagined Movement, Saccades |
| 56 | 32.6 | -6.8 | 56.7 | PrG | Vision.Shape, Motor Learning, Interoception | Anti-Saccades, Drawing, Observation, Imagined Movement, Finger Tapping |
| 61 | -51.9 | 0.3 | 7.5 | PrG | Execution, | Finger Tapping, |

| | | | | | | |
|---|---|---|---|---|---|---|
| | | | | | Perception.Somesthesis .Pain | Pain.Monitor/Discrimination, Isometric Force |
| 62 | 54.3 | 4.5 | 8.4 | PrG | Execution, Perception.Somesthesis .Pain | Chewing/Swallowing, Flexion/Extension, Pain.Monitor/Discrimination, Isometric Force |
| 65 | -8.0 | -38.0 | 57.5 | PCL | - | - |
| 125 | -16.3 | -60.4 | 62.5 | SPL | Imagination, Memory.Working, Execution | Imagined Movement, Mental Rotation, Oddball Discrimination, Drawing, Spatial/Location Discrimination, Saccades |
| 126 | 19.3 | -56.9 | 65.2 | SPL | Cognition.Soma, Vision.Shape, Imagination | Imagined Movement, Action Observation, Saccades, Mental Rotation, Spatial/Location Discrimination |
| 127 | -15.4 | -70.7 | 51.0 | SPL | Attention, Execution, Preparation, Reasoning | Mental Rotation, Counting/Calculation, Anti-Saccades, Imagined Objects/Scenes |
| 128 | 18.7 | -68.9 | 53.3 | SPL | Attention, Execution, Vision.Shape, Memory.Working | Grasping, n-back, Anti-Saccades, Counting/Calculation, Pointing |
| 129 | -33.1 | -47.0 | 49.9 | SPL | Vision.Shape, Cognition.Soma, Execution, Attention | Action Observation, Visual Pursuit/Tracking, Mental Rotation, Pointing, Saccades, Imagined Movement |
| 130 | 34.9 | -41.9 | 53.9 | SPL | Imagination, Cognition.Space, Vision.Motion | Flanker Test, Finger Tapping, Grasping, Drawing |
| 133 | -27.5 | -58.7 | 53.7 | SPL | Cognition.Space, Attention, Reasoning, Memory.Working | Counting/Calculation, Anti-Saccades, Visual Distractor/Visual Attention, Mental Rotation, Saccades |
| 134 | 31.0 | -53.6 | 53.5 | SPL | Interoception.Sexuality, Reasoning, Vision.Motion | Delayed Match To Sample, Orthographic.Discrimination, Saccades, Deduce Reasoning |
| 137 | -37.8 | -61.4 | 46.4 | IPL | Cognition.Space, Reasoning, Memory.Working | n-back, Wisconsin Card Sorting Test, Mental Rotation, Counting/Calculation |
| 138 | 39.7 | -64.8 | 44.0 | IPL | Cognition, Attention, Cognition.Space, Reasoing | Mental Rotation, n-back, Wisconsin Card Sorting Test, Counting/Calculation |
| 139 | -51.4 | -33.2 | 41.4 | IPL | Memory.Working, Attention, Observation | Counting/Calculation, Mental Rotation, Imagined Movement, Finger Tapping |
| 140 | 47.4 | -35.1 | 45.5 | IPL | Memory.Working, Cognition.Time, Perception.Somesthesis | Mental Rotation, Tactile Monitor/Discrimination, Sequence Recall/Learning, Drawing |
| 142 | 57.4 | -43.5 | 38.8 | IPL | Interoception.Blader, Attention, Vision.Motion, Execution | Pain.Monitor/Discrimination, Go/No-Go |

| | | | | | | |
|---|---|---|---|---|---|---|
| 147 | -4.2 | -63.8 | 50.5 | PCun | Vision.Motion, Memory.Working | Visual Distractor/Visual Attention, Imagined Objects/Scenes |
| 148 | 6.6 | -64.6 | 51.3 | PCun | Vision.Motion, Cognition.Space | n-back, Anti-Saccades |
| 150 | 7.5 | -46.6 | 58.5 | PCun | Social Cognition, Cognition.Space | n-back, Deception Task, Spatial/Location Discrimination |
| 159 | -45.6 | -30.2 | 50.3 | PoG | Execution, Perception.Somesthesis | Flexion/Extension, Finger Tapping, Grasping, Tactile Monitor/Discrimination |
| 166 | 33.1 | 14.6 | -12.6 | INS | Gustation, Inhibition | Cued Explicit.Recognition, Go/No-Go, Eating/Drinking |
| 167 | -33.8 | 18.0 | 1.9 | INS | - | - |
| 168 | 36.2 | 18.9 | 0.9 | INS | Inhibition, Cognition, Perception.Somesthesis .Pain | Sternberg Test, Reward Task, Pain Monitor/Discrimination |
| 169 | -38.1 | -3.6 | -9.2 | INS | Perception.Somesthesis , Emotion.Disgust, Perception.Somesthesis .Pain | Acupuncture, Pain Monitor/Discrimination |
| 170 | 39.0 | -1.7 | -9.1 | INS | Emotion.Disgust, Perception.Olfaction | Olfactory Monitor/Discrimination, Pain Monitor/Discrimination |
| 173 | -37.7 | 5.1 | 4.4 | INS | Perception.Somesthesis .Pain, Gustation | Pain Monitor/Discrimination |
| 174 | 38.2 | 6.0 | 4.3 | INS | Perception.Somesthesis .Pain | Pain Monitor/Discrimination, Eating/Drinking, Chewing/Swallowing |
| 180 | 4.8 | 27.7 | 27.7 | CG | Perception.Olfaction, Attention, Inhibition | Olfactory Monitor/Discrimination, Reward Task, Stroop Task, Anti-Saccades |
| 183 | -4.2 | 6.2 | 37.9 | CG | Preparation, Execution, Perception.Somesthesis .Pain | Finger Tapping, Pain Monitor/Discrimination, Pointing |
| 184 | 4.5 | 6.2 | 38.4 | CG | Perception.Somesthesis , Execution, Perception.Somesthesis . Pain | Tactile Monitor/Discrimination, Pain Monitor/Discrimination, Chewing/Swallowing |
| 185 | -7.1 | -22.7 | 40.7 | CG | - | Pointing |
| 186 | 6.4 | -20.8 | 40.6 | CG | Emotion | Reward Task, Pain Monitor/Discrimination |
| 201 | -45.6 | -73.7 | 2.5 | LOcC | Cognition.Space, Vision, Interoception.Sexuality | Visual Distractor/Visual Attention, Action Observation, Subjective Emotional Picture Discrimination, Visual Pursuit/Tracking |

*SFG*, superior frontal gyrus; *MFG*, middle frontal gyrus; *OrG*, orbital gyrus; *PrG*, precentral gyrus; *PCL*, paracentral lobule; *SPL*, superior parietal lobule; *IPL*, inferior parietal lobule; *PCun*, precuneus; *PoG*, postcentral gyrus; *INS*, insula cortex; *CG*, cingulate gyrus; *LOcC*, lateral occipital cortex. The ID is the number of the parcel in the Brainnetome atlas (Fan et al., 2016). Here we only summarized the executive control-related behavioral domains and paradigm classes; the full behavioral domains and paradigm classes for each node are available at http://atlas.brainnetome.org/bnatlas.php.

**Supplementary Table 3. Machine learning-based prediction model accuracies and significance in short-term medication and validation cohorts.**

| | | | Positive FTD | | Negative FTD | |
|---|---|---|---|---|---|---|
| | | | *r* | *p* | *r* | *p* |
| Short-term medication cohort | Language network | State 1 | -0.045 | 0.581 | 0.212 | 0.093 |
| Short-term medication cohort | Language network | State 2 | 0.232 | 0.1 | 0.22 | 0.096 |
| Short-term medication cohort | Language network | State 3 | 0.053 | 0.382 | 0.186 | 0.118 |
| Short-term medication cohort | Language network | State 4 | -0.084 | 0.661 | 0.229 | 0.109 |
| Short-term medication cohort | Language network | State 1+2+3+4 | 0.036 | 0.426 | 0.221 | 0.124 |
| Short-term medication cohort | Executive control network | State 1 | -0.052 | 0.623 | **0.24** | **0.047** |
| Short-term medication cohort | Executive control network | State 2 | 0.168 | 0.119 | **0.458** | **0.001** |
| Short-term medication cohort | Executive control network | State 3 | 0.141 | 0.21 | **0.241** | **0.047** |
| Short-term medication cohort | Executive control network | State 4 | 0.194 | 0.151 | **0.418** | **0.003** |
| Short-term medication cohort | Executive control network | State 1+2+3+4 | 0.076 | 0.26 | **0.395** | **0.004** |
| Validation cohort | Language network | State 1 | **0.261** | **0.009** | 0.098 | 0.245 |
| Validation cohort | Language network | State 2 | **0.477** | < **0.001** | 0.128 | 0.247 |
| Validation cohort | Language network | State 3 | **0.3** | **0.022** | 0.022 | 0.415 |
| Validation cohort | Language network | State 4 | **0.421** | **0.004** | 0.154 | 0.132 |
| Validation cohort | Language network | State 1+2+3+4 | **0.436** | < **0.001** | -0.11 | 0.808 |
| Validation cohort | Executive control network | State 1 | 0.165 | 0.153 | **0.272** | **0.046** |
| Validation cohort | Executive control network | State 2 | **0.355** | **0.007** | 0.182 | 0.121 |
| Validation cohort | Executive control network | State 3 | 0.201 | 0.096 | 0.027 | 0.467 |
| Validation cohort | Executive control network | State 4 | 0.235 | 0.083 | **0.266** | **0.042** |
| Validation cohort | Executive control network | State 1+2+3+4 | 0.25 | 0.057 | **0.331** | **0.009** |

**Supplementary Table 4. Partial correlations between shared language network differences and PANSS without FTD scores.**

| | | | Region 1 | Region 2 | *r* | *p* |
|---|---|---|---|---|---|---|
| Discovery cohort | State 1 | HC > SZ | LSFG | RITG | -0.220 | 0.04 |
| | | HC < SZ | RSFG | RMTG | 0.223 | 0.037 |
| | State 2 | HC < SZ | RMFG | RpSTS | 0.267 | 0.012 |
| | State 3 | HC > SZ | LIFG | LMFG | -0.256 | 0.016 |
| | | | | RMFG | -0.294 | 0.006 |
| | | | | RIFG | -0.210 | 0.049 |
| | | | LITG | LSFG | -0.242 | 0.023 |
| | | | | RIFG | 0.261 | 0.014 |
| | | | RIFG | RMFG | 0.225 | 0.035 |
| | | HC < SZ | RSTG | RITG | -0.327 | 0.002 |
| | | | | LFuG | -0.364 | < 0.001 |
| | | | | RFuG | -0.222 | 0.038 |
| | | | LSTG | LFuG | -0.256 | 0.016 |
| | State 4 | HC > SZ | LSTG | RSTG | 0.246 | 0.021 |
| Short-term medication cohort | State 1 | HC > SZ | RMTG | LFuG | 0.258 | 0.037 |
| | | | | RFuG | 0.351 | 0.004 |
| | | | LMTG | RFuG | 0.282 | 0.022 |
| | | HC < SZ | LIFG | LSTG | 0.252 | 0.041 |
| | | | | LITG | 0.252 | 0.041 |
| | | | | LpSTS | 0.250 / 0.261 | 0.043 / 0.034 |
| | | | | RpSTS | 0.259 | 0.036 |
| | | | LMFG | LpSTS | 0.333 | 0.006 |
| | | | LIFG | RSTG | 0.270 | 0.029 |
| | | | | LITG | 0.246 | 0.047 |
| | State 2 | HC > SZ | LMTG | LFuG | 0.250 | 0.043 |
| | | HC < SZ | RIFG | RSTG | -0.243 | 0.049 |
| | State 3 | HC > SZ | RIFG | LIFG | -0.260 / -0.258 | 0.035 / 0.037 |
| | | | RITG | LpSTS | -0.353 | 0.004 |
| | | | | LMFG | -0.274 | 0.026 |
| | | | | LIFG | -0.292 / -0.324 | 0.018 / 0.008 |
| | | | LITG | RMFG | 0.322 | 0.008 |
| | | | | LIFG | 0.284 | 0.021 |
| | | | | LMTG | 0.278 | 0.024 |
| Validation cohort | State 1 | HC > SZ | LITG | LSTG | 0.215 / 0.262 | 0.009 / 0.002 |
| | | | | RSTG | 0.193 | 0.02 |
| | | HC < SZ | LIFG | LMTG | 0.169 | 0.042 |
| | | | | LITG | 0.168 | 0.044 |
| | State 2 | HC > SZ | LFuG | LIFG | -0.163 / -0.172 | 0.049 / 0.039 |
| | | | | LITG | -0.165 | 0.047 |
| | | | RFuG | LITG | -0.165 | 0.047 |

| | | | | | | |
|---|---|---|---|---|---|---|
| | | HC < SZ | RMTG | RSTG | 0.206 | 0.013 |
| | | | | LPoG | 0.175 | 0.036 |
| | | | | RPoG | 0.199 | 0.016 |
| | | | LMTG | RSTG | 0.181 | 0.03 |
| | | | | RPoG | -0.196 | 0.018 |
| | State 3 | HC > SZ | LITG | RSTG | 0.169 | 0.042 |
| | | | | LMTG | 0.193 | 0.020 |
| | | | | RMTG | 0.218 | 0.008 |
| | | | | LITG | 0.171 | 0.04 |
| | | | LFuG | LSTG | 0.197 | 0.017 |
| | | | | LMTG | 0.170 / 0.172 | 0.039 / 0.041 |
| | | | | LIPL | 0.224 | 0.007 |
| | | | | RIPL | 0.186 | 0.025 |

**Supplementary Table 5. Partial correlations between shared executive control network differences and PANSS without FTD scores.**

| | | | Region 1 | Region 2 | *r* | *p* |
|---|---|---|---|---|---|---|
| Discovery cohort | State 1 | HC > SZ | LSFG | LMFG | -0.236 | 0.023 |
| | | HC < SZ | LSPL | LCG | -0.281 | 0.006 |
| | State 2 | HC > SZ | RINS | LSPL | 0.211 | 0.044 |
| | | | | RSPL | 0.272 | 0.009 |
| | | | | RIPL | 0.241 | 0.021 |
| | | | | LINS | 0.248 | 0.017 |
| | State 3 | HC > SZ | LCG | RSPL | -0.226 | 0.031 |
| | | HC < SZ | RMFG | RSPL | -0.210 | 0.044 |
| | | | | LSPL | -0.206 | 0.049 |
| | State 4 | HC < SZ | LCG | LMFG | 0.272 | 0.009 |
| | | | | LSPL | -0.226 | 0.03 |
| Short-term medication cohort | State 1 | HC < SZ | LSPL | RPrG | 0.280 | 0.021 |
| | State 2 | HC < SZ | RCG | LSPL | 0.306 / 0.256 | 0.011 / 0.036 |
| | State 3 | HC > SZ | RSPL | LCG | -0.264 | 0.03 |
| | | | | RCG | -0.283 | 0.02 |
| | | HC < SZ | RSFG | LSPL | -0.269 | 0.027 |
| | State 4 | HC > SZ | RINS | RCG | 0.243 | 0.046 |
| | | HC < SZ | LCG | LSPL | -0.257 | 0.035 |
| Validation cohort | State 2 | HC > SZ | RMFG | LSPL | -0.188 | 0.02 |
| | State 3 | HC > SZ | RSFG | RINS | -0.171 | 0.034 |

**Supplementary Table 6. sFC-based prediction model accuracies and significance in short-term medication and validation cohorts.**

| | | Positive FTD | | Negative FTD | |
|---|---|---|---|---|---|
| | | *r* | *p* | *r* | *p* |
| Discovery cohort | Language network | **0.36** | **0.004** | -0.044 | |
| | Executive control network | 0.197 | | 0.114 | |
| Short-term medication cohort | Language network | 0.214 | | 0.011 | |
| | Executive control network | 0.016 | | -0.038 | |
| Validation cohort | Language network | **0.397** | **0.001** | 0.095 | |
| | Executive control network | 0.19 | | 0.159 | |
